# Dynamic construction of subjective time through statistical learning of event structure

**DOI:** 10.64898/2026.04.23.720320

**Authors:** Qiyuan Zeng, Darinka Trübutschek, Nicholas B. Turk-Browne, Lucia Melloni

## Abstract

The perception of time is elastic, often deviating from physical intervals depending on how experience is structured. Yet, what determines how subjective time is constructed remains debated. Here, we tested whether perceived ongoing time is actively constructed from learned event representations rather than a dedicated internal clock. Using a novel pause-adjustment task across three statistical learning experiments, we measured temporal distortions during continuous listening to structured versus unstructured syllable streams. The presence of event structure systematically warped time: pauses were perceived as longer between pseudowords and shorter within pseudowords. This bidirectional temporal warping emerged online and remained stable across pause durations. Enriching these events with semantic meaning eliminated boundary-related dilation while preserving within-event compression.

Moreover, physiological tracking of event structure, indexed by pupil dynamics, dissociated from the magnitude of temporal warping. These findings show that subjective time is constructed from hierarchical event representations and depends not only on where events are segmented, but also on how they are represented.

## Introduction

When listening to a language we do not understand, speech can feel strikingly fast. Once that same language becomes familiar, the very same acoustic input appears to slow down. Crucially, this shift occurs without any change in physical timing, suggesting that subjective time depends on how experience is parsed and understood rather than on the stimulus itself.

This intuition is supported empirically. Speech in unfamiliar languages is perceived as faster than acoustically matched speech in familiar languages, even when objective rate is held constant (Bosker & Reinisch, 2015). Manipulations of cognitive load or contextual speech rate similarly bias perceived timing (Bosker et al., 2017), sometimes causing entire words to appear or disappear (Dilley & Pitt, 2010).

Together, these findings point to a close relationship between how continuous input is segmented, how it is interpreted, and how time is experienced.

Classical models of time perception distinguish between prospective and retrospective timing, proposing that ongoing (prospective) experience reflects the output of an internal clock modulated by attention or arousal (Block & Zakay, 1997; Grondin, 2010; Tsao et al., 2022). In contrast, recent accounts suggest that subjective time may emerge from changes in perceptual and mnemonic representations themselves, which could be instantiated by neural population trajectories across the brain (Tsao et al., 2022) without requiring a dedicated timing mechanism. For example, human-like time estimates can arise from accumulating representational change in perceptual classification networks (Roseboom et al., 2019), and moment-to-moment biases in duration judgments can be predicted from sensory cortical dynamics during naturalistic viewing (Sherman et al., 2022). These findings support a constructive view in which time is not read out from a clock, but assembled from the structure of ongoing experience.

A central process that organizes experience is event segmentation—the parsing of continuous sensory input into discrete units such as words, actions, or episodes (Kurby & Zacks, 2008; Zacks et al., 2007). Event segmentation operates online during perception and supports prediction, comprehension and memory (Kurby & Zacks, 2008; Zacks et al., 2007). Importantly, events are organized hierarchically, with lower-level elements nested within higher-level structures (Baldassano et al., 2017). If subjective time depends on how experience is structured, then event segmentation provides a natural candidate mechanism for shaping perceived duration.

Speech provides a clear example of hierarchical event organization. Frequency-tagging studies show that neural activity tracks linguistic structure simultaneously at multiple levels, including syllables, words, phrases, and sentences (Ding et al., 2016, 2017). This hierarchical tracking reflects the brain’s active parsing of continuous input into nested event representations in real time. Such parsing is not unique to language but arises from domain-general statistical learning mechanisms. Even in the absence of meaning, observers rapidly extract temporal regularities and group elements into higher-order units based on transitional probabilities (Aslin & Saffran, 2025; Frost et al., 2019; Isbilen & Christiansen, 2022).

Neural evidence indicates that these statistically defined events are represented across cortex and hippocampus (Henin et al., 2021; Zhou & Turk-Browne, 2025). However, it remains unclear whether and how this hierarchical organization and its underlying representations directly shape the online experience of time.

One possibility is that subjective time emerges from computations operating at multiple levels of event representation. At shorter timescales, prediction dynamics based on transitional probabilities, supported by early sensory and sensory-associative areas e.g., primary auditory and primary visual cortices (Aslin & Saffran, 2025; Batterink & Paller, 2017; Henin et al., 2021), may influence perceived duration, with unexpected or low-probability events experienced as longer (Pariyadath & Eagleman, 2007; Tse et al., 2004). At longer timescales, events are built from clusters of regularly co-occurring elements which become increasingly integrated and, more abstract, supporting chunked, and more holistic representations (Dehaene et al., 2015; Maheu et al., 2019). These representations are supported by medial temporal and prefrontal systems that integrate information across time (Baldassano et al., 2017; Schapiro et al., 2013, 2016; Zhou & Turk-Browne, 2025). Changes in representational structure at these different levels may therefore exert distinct, and potentially opposing, influences on subjective time.

However, direct empirical links between event segmentation, representational format, and perceived duration remain mixed. Studies report both dilation and compression of time around event boundaries (Goh et al., 2025; Ongchoco et al., 2023; Wen et al., 2026), depending on task demands (Zacks et al., 2011) and whether judgments are prospective or retrospective (Block & Zakay, 1997). This variability highlights a critical gap: it remains unclear how event structure shapes subjective time during ongoing perception, rather than in retrospective reconstruction.

Here, we propose that subjective time is actively constructed from learned event representations during ongoing perception, and that this construction depends both on where events are segmented and on how they are internally represented. To test this framework, we introduce a novel online pause-adjustment paradigm that captures temporal distortions as they unfold. Participants continuously adjust brief pauses embedded within or between learned events while listening to ongoing streams, providing a direct measure of subjective time during perception rather than after the fact.

Across three experiments, we combined the pause-adjustment task with a statistical learning paradigm to examine whether and how events structure shapes subjective time. In Experiments 1 and 2, we tested whether learned event structure is sufficient to induce temporal warping during ongoing perception. We found that pauses at event boundaries were perceived as longer (temporal dilation), whereas pauses within events were perceived as shorter (temporal compression), revealing a bidirectional distortion of time.

Experiment 2 further showed that physiological tracking of event structure, indexed by pupil dynamics, dissociates from the magnitude of temporal warping. In Experiment 3, we asked whether altering the representational format of events changes this temporal distortion. When events were enriched with semantic meaning, boundary-related dilation disappeared while within-event compression remained, indicating that how events are represented critically shapes subjective time.

Together, these experiments address a simple but fundamental question: how does the organization of experience give rise to the subjective flow of time? By treating time perception as an emergent property of event construction, this work bridges research on statistical learning, event segmentation, and temporal experience, and provides a new psychophysical framework for studying time as it is lived rather than remembered.

## Results

### Experiment 1: Event segmentation warps subjective time during ongoing perception

Figure 1 summarizes the design and behavioral results of Experiment 1, which tested whether event structure in continuous auditory input is sufficient to distort online subjective time perception.

**Figure 1.**
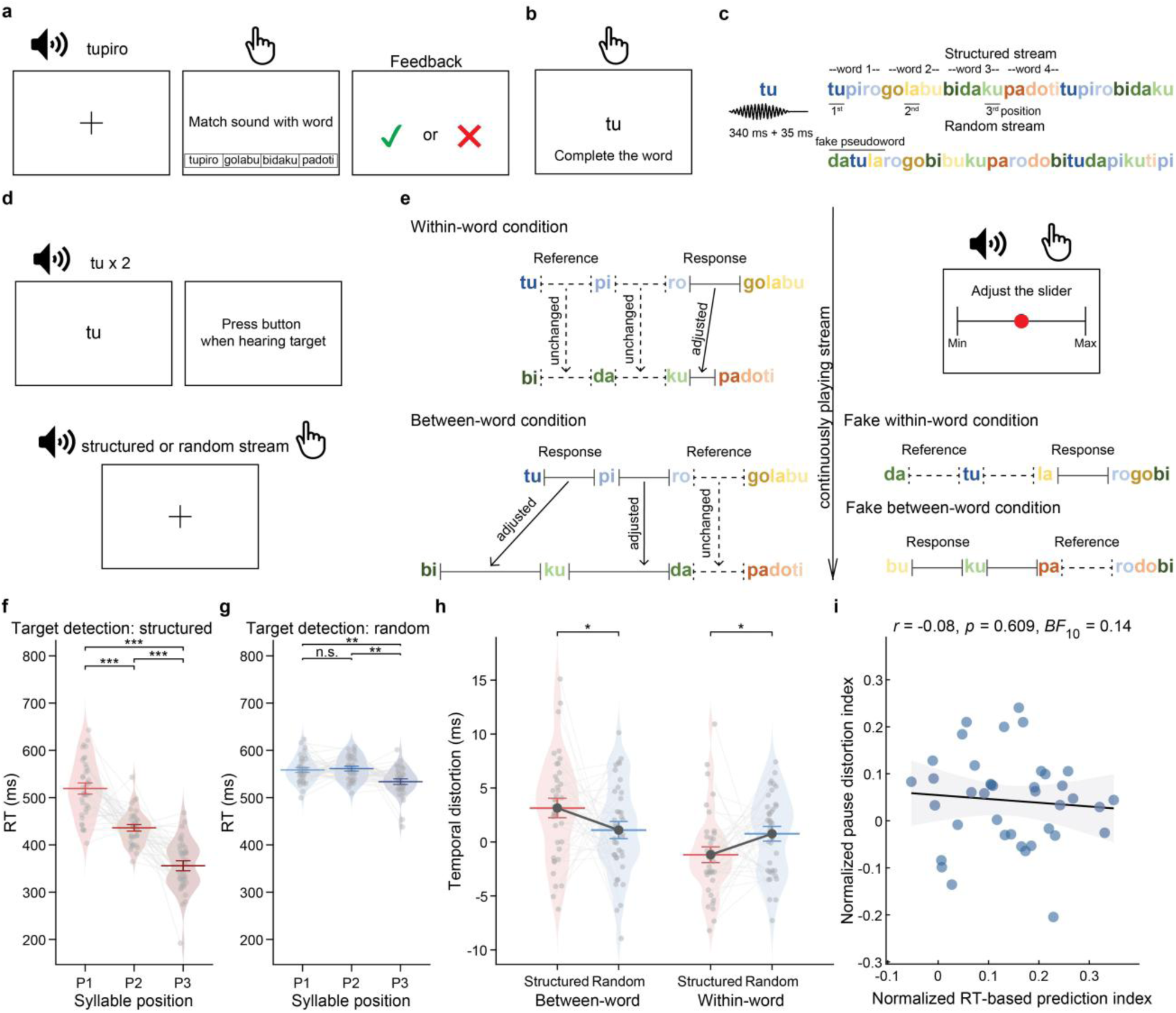
Experimental design and behavioral results of Experiment 1. **a**, Word-matching task. Participants (N = 39) learned four trisyllabic pseudowords by matching auditory presentations to written options. **b**, Stem-completion task. Encoding and retention of pseudoword structure were assessed throughout the experiment using a stem-completion task (12 trials, one per syllable). **c**, Auditory streams. Continuous streams consisted of twelve consonant-vowel syllables (340 ms sound + 35 ms silence). In the structured stream, syllables formed four pseudowords (tupiro, golabu, bidaku, padoti) with transitional probabilities (TPs) of 1.0 within words and 1/3 between words (no back-to-back repetitions). In the random stream, the same syllables were presented in a pseudo-random order (TP = 1/11), eliminating word-level structure. **d**, Target-detection task. Participants detected predefined syllable targets during continuous structured or random streams. **e**, Online pause-adjustment task. While listening continuously, participants adjusted silent pauses using a slider. In the within-word condition, between-word pauses were adjusted to match a fixed within-word reference; in the between-word condition, within-word pauses were adjusted to match a fixed between-word reference. By adjusting to the reference of interest for the condition, the responses indicate the perceived duration of the reference. The same logic was applied to the random streams to fake pseudowords as a control. **f**, Response times in the target detection task in structured streams. Mean target-detection RTs decreased systematically from the first to the third syllable position, indicating successful statistical learning. **g**, RTs in the target detection task in random streams. Without a significant decrease between RTs for positions 1 and 2, mean target-detection RTs showed no systematic facilitation. **h**, Temporal distortion. Relative to random streams, structured streams produced bidirectional temporal distortion: pauses at event boundaries were perceived as longer, whereas pauses within events were perceived as shorter. A linear mixed-effects model revealed a significant interaction between stream type and condition. **i**, Dissociation between learning and time distortion. No significant correlation was observed between the normalized RT-based prediction index and the normalized pause-distortion index. Individual scatter points and violin distributions represent normalized data with between-subject variance removed (panel f-h). Error bars denote Morey-corrected SEM. Shaded regions indicate 95% confidence intervals. Significance was assessed using Holm-corrected post hoc tests following the linear mixed-effects model for temporal distortion analysis (panel h). Significance was assessed using Holm-corrected post hoc tests following ANOVA for RTs analysis (panel f and g), ***P < 0.001; ** P < 0.01; *P < 0.05; N.S., not significant.

Participants (N = 39; mean age = 32.6 ± 8.7 years; LexTALE score = 86.3 ± 8.7; 23 female; 36 right-handed) first learned four trisyllabic pseudowords (tupiro, golabu, bidaku, padoti; Fig. 1a-b). They were then exposed to continuous syllable streams (Fig. 1c) in which these units were either preserved (structured) or disrupted (random), while performing a target-detection task (Fig. 1d), followed by a continuous pause-adjustment task (Fig. 1e).

Memory performance immediately after training was high (mean accuracy: 93.6%), confirming successful encoding of the event units. Although performance declined modestly after exposure to the streams (*t*(38) = 2.551, *p* = 0.015, Cohen’s *d_z_* = 0.47, CI: [1.46, 12.65]), it remained well above chance (≈86%), indicating that the learned event representations were preserved throughout the experiment (Supplementary Fig. 1a). This explicit pre-training, although typically omitted in statistical learning studies, ensured that participants entered the main task with stable event representations, allowing us to isolate their impact on perception.

We next verified that participants segmented the continuous input into event units during listening. Response times (RTs) in the target-detection task revealed a robust interaction between stream type and syllable position (Greenhouse-Geisser correction *F*(1.59, 60.54) = 39.04, *p* < 0.001, partial η² = 0.51). In structured streams, RTs decreased progressively from the first to the third syllable (Greenhouse-Geisser correction *F*(1.56, 59.38) = 66.97, *p* < 0.001, partial η² = 0.64; all pairwise comparisons *p*s < 0.001; Fig. 1f), reflecting increasing predictability within learned units, which is a hallmark of sensitivity to transitional probabilities (Batterink, Reber, Neville, et al., 2015; Batterink, Reber, & Paller, 2015; Batterink & Paller, 2017). In contrast, although random streams showed a weaker effect of position (*F*(2, 76) = 7.80, *p* < 0.001, partial η² = 0.17), they lacked this sequential facilitation, with no difference between the first and second syllables (*t*(38) =-0.409, *p* = 0.685, Cohen’s *d_z_* = 0.07, CI: [-20.24, 14.56], *BF*_10_ = 0.19; Fig. 1g). Accordingly, the RT-based prediction index was significantly larger for structured than random streams (*t*(38) = 7.143, *p* < 0.001, Cohen’s *d_z_* = 1.14, CI: [0.10, 0.17]). These results confirm that participants actively parsed the continuous stream into event units.

Having established online segmentation, we next asked whether event structure alters subjective time perception. Participants performed a continuous pause-adjustment task (Fig. 1e), in which they adjusted the duration of brief silent pauses occurring within or between pseudowords while listening to the ongoing stream. Critically, identical procedures were applied to structured and random streams, except that the latter lacked a transitional structure, providing a stringent control.

A linear mixed-effects model confirmed a significant interaction between stream type (structured vs. random) and pause position (within-vs. between-word; *β* =-0.004, *SE* = 0.001, *t*(9942) =-3.175, *p* = 0.002, CI: [-0.006,-0.002]). In structured streams, pauses at event boundaries were perceived as longer than pauses within events (Estimate= 0.004, *t*(9942) = 4.875, *p* < 0.001, CI: [0.003, 0.006]), whereas no such difference was observed in random streams (Estimate= 0.0003, *t*(9942) = 0.385, *p* = 0.701, CI: [-0.001, 0.002], *BF*_10_ = 0.184). Thus, identical physical intervals were experienced differently depending on their position within the learned event structure.

Decomposing this interaction revealed a striking bidirectional distortion of subjective time. Relative to random streams, structured streams produced a temporal dilation at event boundaries, with between-word pauses perceived as longer (Estimate= 0.002, *t*(9942) = 2.300, *p* = 0.022, CI: [0.0003, 0.004]), and a temporal compression within events, with within-word pauses perceived as shorter (Estimate=-0.002, *t*(9942) =-2.190, *p* = 0.029, CI: [-0.004,-0.0002]; Fig. 1h).

This distortion was robust across reference durations (35, 55, 75, and 95 ms). Although absolute pause adjustments scaled with physical duration (main effect of reference level: *β* = 1.077, *SE* = 0.028, *p* < 0.001, CI: [1.022, 1.132]), the relative distortion between within-and between-event pauses remained stable. There was no evidence for a three-way interaction between stream type, pause position, and reference level (*β* =-0.019, *SE* = 0.056, *t*(9938) =-0.335, *p* = 0.737, CI: [-0.129, 0.091]), indicating that temporal warping is preserved across time scales. Consistent with this, the ratio of perceived between-word to within-word pauses was significantly greater than 1 at every reference level (all *p*s < 0.05; Supplementary Fig. 1b).

Finally, we tested whether temporal distortion reflects the strength of statistical learning. Although participants showed robust RTs facilitation and temporal warping at the group level, these measures were not correlated across individuals (*r* =-0.085, *p* = 0.609, CI: [-0.390, 0.237], *BF*_10_ = 0.142 < 1/3; Fig. 1i). Participants with stronger sensitivity to transitional probabilities did not exhibit larger distortions of subjective time. This dissociation indicates that, while event segmentation is necessary for temporal warping to emerge, the magnitude of the distortion is not a simple byproduct of predictive facilitation.

Together, these results demonstrate that learning event structure is sufficient to reshape subjective time during ongoing perception. Event boundaries selectively expand perceived duration, whereas coherent event units compress it, yielding a bidirectional warping of time that emerges only when structured events are present. These findings provide direct evidence that subjective time is actively constructed from event segmentation as experience unfolds.

### Experiment 2: Replication of temporal warping and physiological tracking of event structure

Experiment 2 pursued two complementary goals: to test whether the temporal distortions observed in Experiment 1 replicate under modified task conditions, and to examine whether physiological signals, specifically pupil dynamics (Fig. 2a), track event structure and relate to behavioral measures of learning and time perception. Participants (N = 37; mean age = 29.7 ± 6.5 years; LexTALE score = 84.5 ± 8.4; 15 female; 36 right-handed) completed the same core tasks as in Experiment 1, including pseudoword learning, target detection, and pause adjustment, with target detection and pause adjustment trials interleaved within blocks.

**Figure 2.**
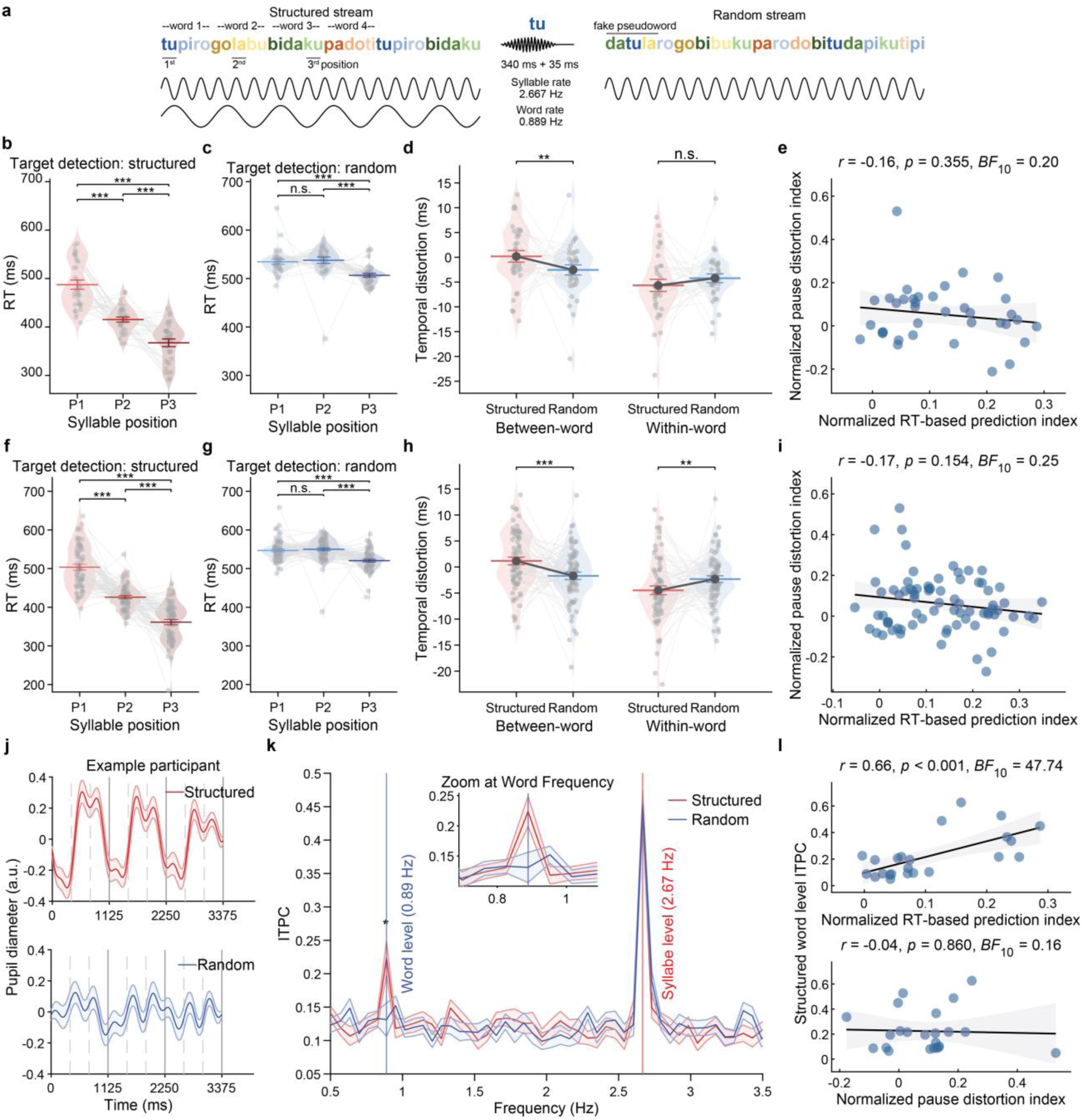
**Replication of temporal warping and physiological tracking of event structure**. **a**, Eye-tracking and pupil data were collected to quantify entrainment to the statistical structure of the auditory streams and to obtain an individualized physiological measure of learning. Auditory streams were extended to 81 s during the target detection task to allow for frequency-domain analysis of pupil dynamics. Syllables were presented at a rate of 2.667 Hz and thus the trisyllabic words appeared at a third of the rate, 0.889 Hz. **b-e**, Behavioral replication of Experiment 1 in Experiment 2 (N = 37). **b**, Target detection RTs significantly decreased across syllable positions in structured streams, replicating the statistical learning effect. **c**, As a control, target detection RTs did not show a sequential decrease in random streams. **d**, Real-time pause adjustment revealed a significant interaction between stream and condition, driven by longer perceived pauses between-word, and a trend for shorter pauses within-word. **e**, Scatter plot showing no significant correlation between the normalized RT-based prediction index and the normalized pause distortion index. **f-i**, Combined behavioral analysis pooling Experiments 1 and 2 (N = 76). **f**, Pooled results confirmed robust position-dependent RT facilitation in structured streams. **g**, Such RT facilitation was absent in random streams for the combined cohort. **h**, Combined analysis confirmed bidirectional warping with temporal dilation at event boundaries and temporal compression within events. **i**, Correlation analysis for the combined dataset confirmed the dissociation between statistical prediction and time distortion. **j-l**, Behavior and eyetracking data analysis with pupil data from selected participants (N=23). **j**, Pupil entrainment was evident at the single-subject level. Pupil diameter dynamics for one example participant during exposure to structured stream (upper panel, red) showed clear modulation at the syllable rate (375 ms, grey vertical dash line) and word rate (1125ms, grey vertical solid line). The predominant frequency in the random streams (lower panel, blue) was at syllable rate. **k**, Inter-trial phase coherence (ITPC) of pupil diameter. Structured streams elicited significant entrainment at both syllable (2.67 Hz) and word (0.89 Hz) frequencies, whereas random streams exhibited only syllable-level tracking (2.67 Hz). **l**, Word-level pupil ITPC significantly correlated with the normalized RT-based prediction index, identifying pupil entrainment as a physiological marker of sensitivity to transitional probabilities in statistical learning (upper panel). Word-level pupil ITPC did not correlate with the normalized pause-distortion index, suggesting distinct cognitive mechanisms behind physiological tracking (and RT facilitation) versus temporal warping (lower panel). Individual scatter points and violin distributions represent normalized data with between-subject variance removed (panel b-d, f-h). Shaded areas in correlation plots represent the 95% confidence interval (CI). Shaded areas in the ITPC spectra and error bars in panels (b-d, f-h, k) represent Morey-corrected SEM, with normal SEM in panel (j). Significance for panels (d, h) were determined via Holm-corrected post-hoc simple effects from the LMM. Significance was assessed using Holm-corrected post hoc tests following ANOVA analysis (panel b, c, f, g and k). ***P < 0.001; **P < 0.01; N.S., not significant.

Participants successfully learned and retained the pseudowords throughout the experiment. Memory performance remained stable from the initial test (mean = 86.3%) to the mid-experiment test (87.8%; *t*(36) =-0.468, *p* = 0.643, *BF*_10_ = 0.196, CI: [-8.409, 5.256], Cohen’s *d_z_* = 0.10; Comparisons with Experiment 1 were further showed in Supplementary Fig. 2a), indicating robust retention of event representations. Statistical learning during continuous listening replicated the pattern observed in Experiment 1. Response times showed a strong interaction between stream type and syllable position (Greenhouse-Geisser corrected *F*(1.68, 60.50) = 24.92, *p* < 0.001, partial η² = 0.41), with structured streams eliciting the characteristic sequential facilitation across syllable positions (Greenhouse-Geisser corrected *F*(1.45, 52.19) = 60.29, *p* < 0.001, partial η² = 0.63; all adjacent comparisons *p*s < 0.001; Fig. 2b), a pattern absent in random streams (without significant comparison between first and second position in simple analysis: t(36)=-0.361, p = 0.720, CI: [-26.50, 19.80], Conhen’s *d_z_* = 0.06, *BF*_10_ = 0.19; Fig. 2c). The magnitude of this effect did not differ from Experiment 1 (*t*(74) = 1.049, *p* = 0.298, CI: [-0.02, 0.07], Cohen’s *d* = 0.24, *BF*_10_ = 0.382; Supplementary Fig. 2b), indicating comparable levels of statistical learning across experiments.

Crucially, the temporal distortion observed in Experiment 1 replicated. Pause adjustments revealed a significant interaction between stream type and pause position (*β* =-0.004, *SE* = 0.001*, t*(7064) =-2.862, *p* = 0.004, CI: [-0.007,-0.001]). In structured streams, pauses at event boundaries were perceived as longer than pauses within events (Estimate = 0.006, *t*(7064) = 5.670, *p* < 0.001, CI: [0.004, 0.008]), whereas no reliable difference was observed in random streams (Estimate = 0.002, *t*(7064) = 1.622, *p* = 0.105, CI: [-0.0003, 0.004], *BF*_10_ = 1.66). This effect was primarily driven by a dilation at event boundaries, with between-word pauses perceived as longer in structured than in random streams (Estimate = 0.003, *t*(7064) = 2.643, *p* = 0.008, CI: [0.001, 0.005]), while within-event compression was not significant in this experiment (Estimate =-0.001, *t*(7064) =-1.405, *p* = 0.160, CI: [-0.003, 0.001], *BF*_10_ = 0.256; Fig. 2d). Thus, the key signature of event-induced temporal warping, particularly boundary-related dilation, was robust across experimental conditions.

The temporal distortion generalized across the two reference pause durations tested in Experiment 2 (55 and 75 ms; *β* =-0.054, *SE* = 0.146, *t*(7060) =-0.372, *p* = 0.710, CI: [-0.34, 0.23]). The ratio of perceived between-word to within-word pauses was significantly greater than 1 at both reference levels (55 ms: *t*(36) = 3.900, *p*_FDR_ < 0.001; 75 ms: *t*(36) = 3.820, *p_FDR_* < 0.001; Supplementary Fig. 2c), indicating that the effect was not tied to a specific time scale.

As in Experiment 1, temporal distortion was not explained by individual differences in statistical learning. The magnitude of pause distortion was not correlated with the RT-based prediction index (*r* =-0.157, *p* = 0.355, CI: [-0.46, 0.18], *BF*_10_ = 0.20 < 1/3; Fig. 2e), indicating that temporal warping is not a simple downstream consequence of predictive facilitation.

To more sensitively test the relationship between temporal distortion and statistical learning, we pooled data from Experiments 1 and 2 (N = 76), thereby increasing statistical power for individual-difference analyses. The combined dataset confirmed robust statistical learning as shown by RTs facilitation in structured streams (Greenhouse-Geisser corrected *F*(1.52, 114.03) = 121.58, *p* < 0.001, partial η² = 0.62; Fig. 2f) with random streams as control (Fig. 2g), and reliable temporal distortion effects (within-vs. between-word; *β* =-0.004, *t*(12020) =-4.394, *p* < 0.001, CI: [-0.007,-0.003]), with pauses between pseudowords perceived as significantly longer than pauses within pseudowords (Estimate = 0.006, *t*(12017) = 7.27, *p* < 0.001, CI: [0.004, 0.007]) in structured streams. No such difference was observed in the random streams (Estimate = 0.0008, *t*(12017) = 1.056, *p* = 0.291, CI: [-0.0007, 0.0024], *BF*_10_ = 0.180 < 1/3). And more importantly, the bidirectional distortion was also evident in this combined cohort, with between-word pauses perceived as longer (Estimate = 0.003, *t*(12017) = 3.630, *p* < 0.001, CI: [0.001, 0.004]), and within-word pauses perceived as shorter (Estimate =-0.002, *t*(12017) =-2.584, *p* = 0.010, CI: [-0.004,-0.0005]; Fig. 2h).

These results ensured sufficient variability in both measures to assess their relationship. Critically, even with increased statistical power, temporal distortion remained uncorrelated with the RT-based prediction index (*r* =-0.165, *p* = 0.154, CI: [-0.377, 0.063], *BF*_10_ = 0.249 < 1/3; Fig. 2i). Thus, participants who exhibited stronger sensitivity to transitional probabilities did not show larger distortions of subjective time. This null relationship, observed under substantially improved power, provides strong evidence that temporal warping is not a simple downstream consequence of predictive facilitation, but instead reflects a partially dissociable process.

Taken together, these behavioral results demonstrate that event segmentation robustly and bidirectional reshapes subjective time perception, and that this effect is dissociable from standard behavioral indices of learning of transitional probabilities in statistical learning. We next turned to physiological measures to ask whether the brain tracks the hierarchical event structure that gives rise to these temporal distortions.

### Pupil dynamics track hierarchical event structure

To assess whether physiological signals reflect learned event structure, we analyzed pupil diameter during the target detection task in a subset of participants with high-quality eye-tracking data (N = 23; see Methods for participants selection; see Supplementary Fig. 2d-g for replication of behavioral results). In structured streams, inter-trial phase coherence (ITPC; comparable results were obtained on power spectra, see Supplementary Fig. 2h) revealed significant spectral peaks compared to permutated bassline at both the syllable frequency (2.667 Hz, *t*(22) = 7.072, *p* < 0.001, CI: [0.23, 0.43], Cohen’s *d_z_* = 2.03) and the word frequency (0.889 Hz, *t*(22) = 3.078, *p* = 0.006, CI: [0.03, 0.18], Cohen’s *d_z_* = 0.88), whereas in random streams, only syllable-level tracking was observed (2.667 Hz, *t*(22) = 7.665, *p* < 0.001, CI: [0.24, 0.43], Cohen’s *d_z_* = 2.21; 0.889 Hz, *t*(22) = 0.879, *p* = 0.390, CI: [-0.02, 0.04], Cohen’s *d_z_* = 0.24, *BF*_10_ = 0.31 < 1/3). A frequency × stream type interaction (*F*(1, 22) = 5.484, *p* = 0.029, partial η^2^ = 0.20) confirmed selective tracking of learned event structure at the word level (word: *t*(22) = 2.630, *p* = 0.015, CI: [0.02, 0.17], Cohen’s *d_z_*= 0.56; syllable: *t*(11) =-0.006. *p* = 0.854, CI: [-0.07. 0.06], Cohen’s *d_z_* = 0.04, *BF*_10_ = 0.22 < 1/3; Fig. 2k).

Crucially, pupil-based measures of event tracking were selectively related to statistical learning but not to temporal distortion. Word-level ITPC in structured streams correlated strongly with the RT-based prediction index (*r* = 0.655, *p* < 0.001, CI: [0.33, 0.84]; Fig. 2l, upper panel), but not with the magnitude of subjective time distortion (*r* =-0.039, *p* = 0.860, CI: [-0.44, 0.38], *BF*_10_ = 0.16 < 1/3; Fig. 2l, lower panel). No correlation was observed in random streams between word-level ITPC and the normalized RT-based prediction index (*r* =-0.083, *p* = 0.707, *BF*_10_ = 0.17 < 1/3). Thus, pupil dynamics provide a physiological marker of sensitivity to transitional probabilities, but do not appear to participate directly in the construction of subjective time.

Together, these results show that event-induced temporal distortion is robust across independent samples and task configurations. Experiment 2 provides physiological evidence that auditory event structure is tracked hierarchically in pupil dynamics, extending frequency-tagging approaches beyond vision (Binda et al., 2025; Schwiedrzik & Sudmann, 2020). Critically, physiological tracking of event structure, as reflected in pupil dynamics, dissociates from subjective time distortion, suggesting that while both depend on learned structure, they reflect partially distinct cognitive mechanisms.

### Experiment 3: Semantic enrichment reshapes temporal distortion

Whereas Experiments 1 and 2 established that statistically learned event structure is sufficient to induce temporal warping, real-world events, particularly in language, are typically imbued with meaning. Such semantic enrichment may increase the internal coherence of event units by binding their constituents into a unified conceptual representation (Dehaene et al., 2015). We therefore asked whether this change in representational format qualitatively reshapes the temporal warping effect.

Participants (N = 53; mean age = 31.8 ± 8.6 years; LexTALE score = 85.4 ± 8.9; 27 female; all right-handed, native German speakers) learned the same trisyllabic pseudowords as in previous experiments, but each was explicitly associated with a semantic referent (Fig. 3a–b). Training continued until participants achieved perfect recall, ensuring that event representations were both statistically predictable and semantically enriched. Participants then completed the same target detection and pause adjustment tasks as in Experiments 1 and 2.

**Figure 3.**
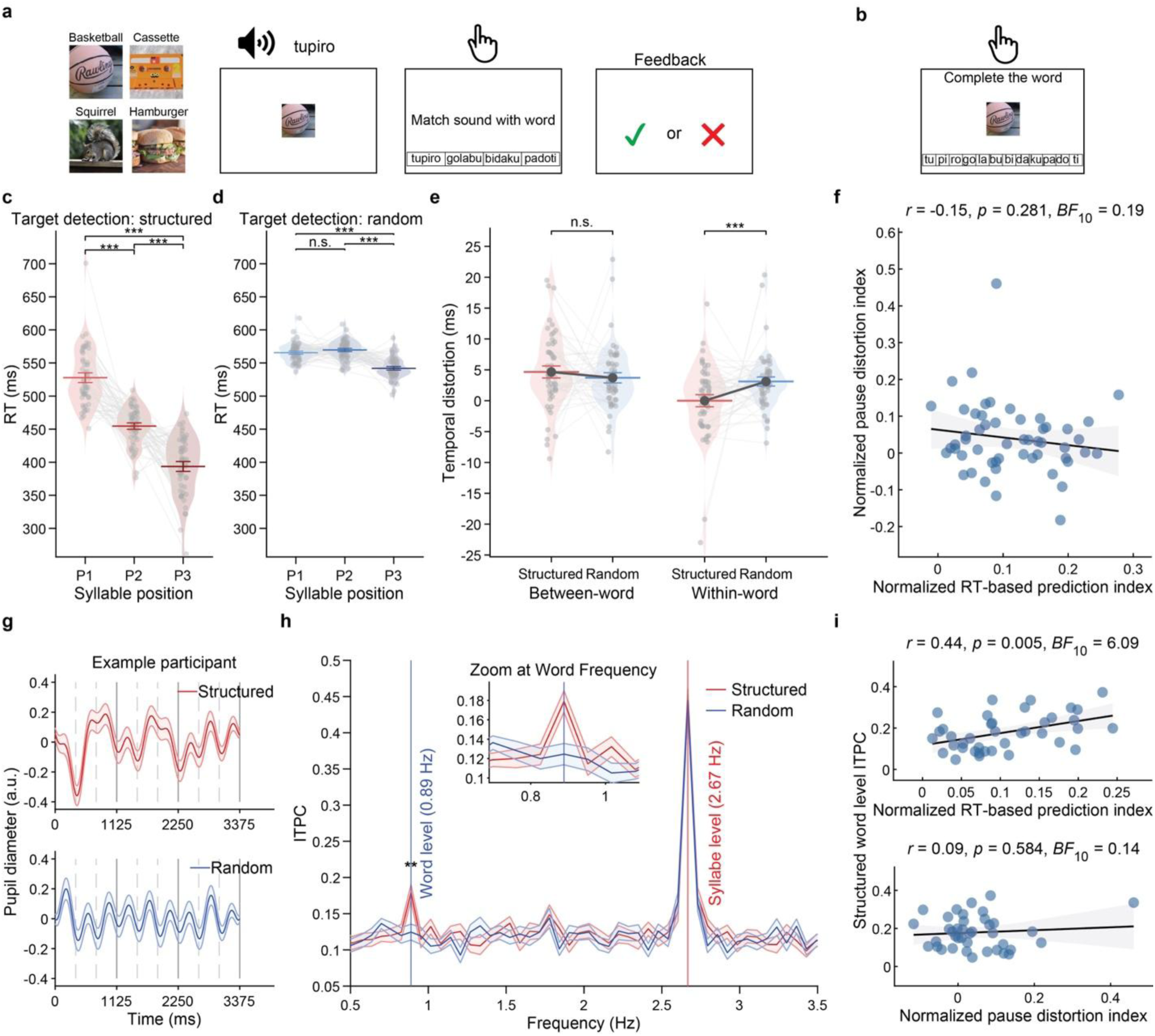
Experiment 3 design and results. **a**, Semantic association paradigm. Participants first learned to associate the four trisyllabic pseudowords (tupiro, golabu, bidaku, padoti) with distinct object categories (e.g., basketball, squirrel, hamburger, cassette). **b**, Semantic memory task. Participants then viewed an image and reconstructed the corresponding pseudoword by selecting its constituent syllables in the correct order. **c-f**, Temporal distortion under semantic enrichment (N=53). **c**, Mean RTs significantly decreased with syllable positions in structured streams. **d**, Such sequential decrease of RTs was absent in random streams. **e**, Semantic enrichment altered the profile of temporal distortion, with no dilation between words but robust compression within words. **f**, No significant correlation was found between the normalized RT-based prediction index and the normalized pause distortion index. **g-i**, Pupil size frequency tagging analysis (N = 39). **g**, Pupil diameter from one sample participant. Pupil diameter tracked syllable (375 ms, grey dash line) and word (1125 ms, grey solid line) rates in structured streams (upper panel, red), with the dominant frequency in random streams (lower panel, blue) at the syllable rate. **h**, Inter-trial phase coherence (ITPC) of pupil diameter revealed clear peaks at both syllable (2.667 Hz) and word (0.889 Hz) frequencies in structured streams, confirming physiological sensitivity to hierarchical structure under semantic enrichment. **i**, Structured word-level pupil ITPC significantly correlated with RT facilitation (upper panel) but not time distortion (lower panel). Individual scatter points and violin distributions represent normalized data with between-subject variance removed (panel c-e). Shaded areas in correlation plots represent the 95% confidence interval (CI). Shaded areas in the ITPC spectra and error bars in panels (c-e, h) represent Morey-corrected SEM. Significance for panel (e) was determined via Holm-corrected post-hoc simple effects from the LMM. Significance was assessed using Holm-corrected post hoc tests following ANOVA analysis (panel c, d and h). ***P < 0.001; **P < 0.01; N.S., not significant.

We explored memory performance across Experiments 1–3 to evaluate whether semantic enrichment supported more persistent event representations. Memory performance was higher in Experiment 3 than in Experiments 1 and 2 combined (*F*(1, 127) = 14.208, *p* < 0.001, partial η ^2^ = 0.06; Supplementary Fig. 3a), indicating more robust event representations. Critically, this enrichment did not alter online segmentation dynamics. Target-detection RTs showed the characteristic positional facilitation in structured streams (Greenhouse-Geisser corrected *F*(1.62, 84.22) = 101.74, *p* < 0.001, partial η² = 0.66), with progressively faster responses across syllables (all *p*s < 0.001; Fig. 3c), but not in the random streams (Fig. 3d), and the magnitude of this effect did not differ from Experiments 1 and 2 (*t*(127) = 1.155, *p* = 0.250, CI: [-0.01, 0.05], Cohen’s *d* = 0.21,*BF*_10_ = 0.35; Supplementary Fig. 3b). Thus, participants successfully segmented the streams, allowing us to isolate how semantic representations alter subjective time.

### Semantic enrichment eliminates boundary dilation and selectively drives within-event compression

Temporal distortion was again observed, as reflected in a significant interaction between stream type and pause position (*β* =-0.004, *SE* = 0.001, *t*(15208) =-3.875, *p* < 0.001, CI: [-0.006,-0.002]). In structured streams, pauses between pseudowords were perceived as longer than pauses within pseudowords (Estimate = 0.005, *t*(15208)= 6.288, *p* <0.001, CI: [0.003, 0.006]), whereas no reliable difference was observed in random streams (Estimate = 0.0006, *t*(15208)= 0.807, *p* = 0.420, CI: [-0.0001, 0.002], *BF*_10_ = 0.21 < 1/3). The overall magnitude of temporal distortion did not differ from Experiments 1 and 2 (*t*(127) = 1.020, *p* > 0.05, CI: [-0.02, 0.07], Cohen’s *d* = 0.18, *BF*_10_ = 0.35; Supplementary Fig. 3c), indicating that semantic enrichment preserved the presence of the effect.

Crucially, however, semantic enrichment qualitatively reshaped the structure of temporal distortion. Decomposition by pause position revealed that, unlike in Experiments 1 and 2, temporal warping was driven selectively by compression within events. Within-word pauses were perceived as shorter in structured than in random streams (Estimate =-0.003, *t*(15208) =-4.208, *p* < 0.001, CI: [-0.005,-0.002]), whereas between-word pauses did not differ reliably (Estimate = 0.001, *t*(15208) = 1.273, *p* = 0.203, CI: [-0.001, 0.002], *BF*_10_ = 0.19 < 1/3; Fig. 3e). Thus, semantic enrichment eliminated the boundary-related dilation observed for purely statistical events while preserving, and even emphasizing, within-event compression. This pattern suggests that semantic binding increases the internal coherence of event representations, pulling their constituent elements together in time while smoothing transitions across event boundaries.

As in the previous experiments, temporal distortion remained dissociable from statistical learning. Individual differences in pause distortion were not correlated with the RT-based prediction index (*r* =-0.151, *p* = 0.281, CI: [-0.40, 0.12] *BF*_10_ = 0.192 < 1/3; Fig. 3f), indicating that the reshaping of subjective time under semantic enrichment does not reflect variation in predictive facilitation.

Together, these results show that semantic enrichment alters the balance of temporal distortion. Whereas purely statistical events induce bidirectional warping, dilating time at boundaries and compressing time within events, semantically enriched events selectively compress perceived time while eliminating boundary-related dilation. Experiment 3 therefore demonstrates that subjective time is shaped not only by where events are segmented, but also by how those events are internally represented. Specifically, semantic enrichment may transform a sequence of syllables into a singular, unitized conceptual object.

This increased coherence pulls the constituents together, thereby intensifying temporal compression within the event while bridging the transition between units, which effectively eliminates the boundary-driven dilation observed for purely statistical structures.

### Semantic enrichment preserves pupil-based frequency tagging of event structure

We next examined whether physiological tracking of event structure is preserved under semantic enrichment. Pupil dynamics were analyzed in a subset of participants with high-quality eye-tracking data (N = 39, see Supplementary Fig. 3d-g for replication of behavioral results). In structured streams, inter-trial phase coherence (ITPC; similar results were obtained for power spectra, see Supplementary Fig. 3h) revealed clear peaks at both the syllable frequency (2.667 Hz) and the word frequency (0.889 Hz) compared to permutated baseline, whereas in random streams, only syllable-level tracking was observed. Specifically in structured streams, ITPC was significantly elevated at both syllable (*t*(38) = 8.278, *p* < 0.001, CI: [0.25, 0.41], Cohen’s *d_z_* = 1.86) and word frequencies (*t*(38) = 4.257, *p* < 0.001, CI: [0.03, 0.08], Cohen’s *d_z_* = 0.91), whereas in random streams only syllable-level ITPC differed from baseline (*t*(38) = 8.233, *p* < 0.001, CI: [0.25, 0.41], Cohen’s *d_z_* = 1.82), with no reliable word-level tracking (*t*(38) = 0.734, *p* = 0.468, CI: [-0.01, 0.02], Cohen’s *d_z_* = 0.16, *BF*_10_ = 0.22 < 1/3). A frequency × stream interaction (*F*(1, 38) = 5.898, *p* = 0.02, partial η^2^= 0.13) confirmed selective tracking of hierarchical event structure, driven by selective phase locking to learned word-level structure in structured vs. random streams (word: *t*(38) = 3.488, *p* = 0.001, CI: [0.02, 0.09], Cohen’s *d_z_* = 0.57; syllable: *t*(38) = 0.212, *p* = 0.833, CI: [-0.03, 0.04], Cohen’s *d_z_*= 0.03, *BF*_10_ = 0.18 < 1/3;Fig.3h).

Crucially, pupil-based measures again dissociated from subjective time. Word-level ITPC in structured streams correlated with the RT-based prediction index (*r* = 0.440, *p* = 0.005, CI: [0.14, 0.66]; Fig. 3i upper panel), but not with temporal distortion (*r* = 0.091, *p* = 0.584, CI: [-0.23, 0.39], *BF*_10_ = 0.14 < 1/3; Fig. 3i lower panel). No correlation was observed in random streams between the word-level ITPC and the normalized RT facilitation index (*r* =-0.029, *p* = 0.860, CI: [-0.34, 0.29], *BF*_10_ = 0.127 < 1/3). Thus, replicating Experiment 2, physiological tracking of event structure reflects sensitivity to transitional probabilities but does not predict subjective time warping.

Together, these results show that semantic enrichment does not merely strengthen event segmentation, but fundamentally alters how event structure shapes subjective time. Whereas purely statistical events induce bidirectional warping, with dilation at boundaries and compression within events, semantically enriched events selectively compress perceived time while eliminating boundary-related dilation. These findings demonstrate that subjective time depends not only on where events are segmented, but also on how those events are represented, with semantic coherence reshaping the temporal structure of experience.

## Discussion

The subjective passage of time is rarely a faithful reflection of physical duration; rather, it is highly malleable, actively constructed from how our ongoing experience is organized. Here we demonstrate that learning the statistical structure of an auditory stream, held constant in its physical timing, can give rise to rapid and systematic distortions of perceived duration online. Across three behavioral experiments, subjective time warped in a lawful manner: when participants learned to segment a continuous syllable stream into pseudowords, pauses at event boundaries were consistently perceived as longer than physically identical pauses within events. Crucially, the specific pattern of this temporal distortion depended on the underlying representational format. When events were defined by statistical regularities, time warped bidirectionally, which was expanding at boundaries and compressing within. However, when these events were enriched with semantic meaning, this distortion pattern shifted qualitatively, eliminating boundary dilation and selectively driving within-event compression. This distortion was expressed online, during continuous listening, using a pause-adjustment procedure that minimized memory demands and anchored responses to ongoing perception, providing direct empirical evidence that subjective time is an actively constructed property of hierarchical event representations.

### Event segmentation distorts subjective time in real time

Classical models of time perception have emphasized an internal clock whose output is passively modulated by attention, arousal, or cognitive load (Block & Zakay, 1997; Grondin, 2010). However, a growing body of work suggests that subjective duration can instead be constructed from changes in perceptual and mnemonic representations themselves, without invoking a centralized pacemaker (Eagleman & Pariyadath, 2009; Roseboom et al., 2019; Sherman et al., 2022). The present findings extend this constructive view by showing that statistical learning, the organization of continuous input into discrete events, reshapes temporal experience as it unfolds.

In Experiments 1 and 2, statistical learning of transitional probabilities was sufficient to induce a bidirectional distortion of subjective time: perceived duration expanded at event boundaries and contracted within events. This pattern cannot be explained by a uniform speeding or slowing of an internal clock. Instead, it suggests that different components of event processing exert opposing influences on temporal experience. Event boundaries, marked by drops in predictability and increases in prediction error, appear to amplify perceived duration, whereas predictable content within events is compressed. Importantly, this dissociation implies that subjective time is not governed by a single scalar mechanism, but by multiple processes operating at different levels of event representation.

This interpretation is consistent with predictive processing accounts in which surprising or low-probability events are experienced as longer than expected ones (Eagleman & Pariyadath, 2009; Pariyadath & Eagleman, 2007). At boundaries between pseudowords, transitional probabilities are low and prediction error is high, plausibly increasing processing demands and expanding perceived time. Conversely, within events, high predictability may promote repetition suppression and efficient encoding, leading to temporal compression (Pariyadath & Eagleman, 2007). However, prediction-based accounts alone are insufficient to explain the full pattern observed here, particularly the persistence of compression and the dissociation from behavioral indices of prediction strength.

### An integrative account: boundary updating and within-event integration

Statistical learning not only sharpens predictions but also reorganizes representations, clustering elements into chunked units or “events” (Henin et al., 2021; Schapiro et al., 2013, 2016). As sequences are learned, neural representations of elements within an event become more similar to each other, particularly in medial temporal lobe structures including the hippocampus (Henin et al., 2021; Schapiro et al., 2012, 2016). In computational terms, this reduces the geometric distance between successive elements within an event.

Recent models of time perception propose that subjective duration reflects the distance traversed through neural state space rather than elapsed physical time (Tsao et al., 2022). From this perspective, within-event compression may arise because hierarchical chunking collapses the constituent sensory elements toward a common representation, shortening the neural distance between the elements. Critically, this mechanism operates independently of local prediction error, providing a principled account of why temporal compression does not scale with behavioral measures of predictability. This account predicts temporal compression within events irrespective of local prediction error and thus a dissociation from response-time facilitation (Schapiro et al., 2013), which primarily indexes predictability rather than representational distance.

Experiment 3 can be interpreted through this representational account. When pseudowords were enriched with semantic meaning, the magnitude of temporal distortion remained comparable to Experiments 1 and 2, but between-event expansion disappeared (while within-event compression persisted). This pattern is difficult to explain solely in terms of statistical learning, as response time measures were indistinguishable across experiments. Instead, semantic enrichment appears to alter how events are represented and accessed. When a pseudoword becomes associated with a semantic concept, it may be less likely to be processed as a decomposable syllable sequence and instead treated as a unified, conjunctive representation. This shift effectively reduces the need for continuous boundary-based updating at the level of syllables, thereby attenuating the conditions that give rise to temporal dilation.

Cognitive and neural models demonstrate that when continuous input is parsed into meaningful linguistic units, the brain dynamically restructures its predictive computations: rather than solely computing local, transitional predictions (Peña et al., 2002), the cognitive system increasingly relies on higher-level, chunk-based mechanism (Kuperberg & Jaeger, 2016). Critically, recent evidence suggests that this chunking mechanism constrains prediction by enhancing anticipatory processing within a unit while compressing prior information into a global context at boundaries (Ding et al., 2025). Under these conditions, boundaries are no longer experienced as points of local surprise but as transitions between coherent, already-integrated representations, reducing their impact on subjective time.

Taken together, the three experiments support an integrative framework in which subjective time emerges from the interaction of two complementary processes: boundary updating and within-event integration.

Event boundaries mark points at which internal models are updated, increasing salience and expanding perceived duration. Within events, predictive stability and representational clustering promote integration, compressing perceived time. When events are defined purely by transitional probabilities, both processes contribute, yielding bidirectional warping. When events are semantically enriched, integration dominates and boundary-related expansion is attenuated. This framework provides a unified account that reconciles prediction-based and representational accounts by assigning them distinct roles within the temporal construction of experience.

Crucially, this account does not posit a dedicated timing mechanism. Instead, perceived duration emerges as a byproduct of how experience is organized into hierarchical event representations—a view consistent with broader proposals that time perception reflects representational dynamics rather than an internal clock (Buzsáki, 2026; Eagleman & Pariyadath, 2009; Tsao et al., 2022). In this sense, time perception may be better understood as an emergent property of cognitive architecture rather than a primary function in its own right.

### Pupil entrainment tracks event learning but not temporal warping

The present work also provides physiological evidence of statistical learning. Using frequency-tagging, pupil diameter entrained not only to syllable-level rhythms but also to word-level structure. This extends prior demonstrations of pupil entrainment to structure from the visual to the auditory domain (Binda et al., 2025; Schwiedrzik & Sudmann, 2020) and complements neural frequency-tagging studies of speech and statistical learning (Ding et al., 2016; Henin et al., 2021).

Importantly, pupil entrainment was tightly linked to behavioral sensitivity to transitional probabilities: individuals with stronger word-level pupil phase coherence showed greater response time facilitation. This supports the view that pupil dynamics index the strength of internal models used for prediction, potentially via locus coeruleus–norepinephrine mechanisms that regulate neural gain and orienting responses (Aston-Jones & Cohen, 2005; Jepma & Nieuwenhuis, 2011).

At the same time, pupil entrainment did not correlate with subjective time distortion. This dissociation suggests that physiological tracking of structure and temporal warping reflect distinct facets of event processing. One possibility, consistent with a dual-system account of statistical learning (Zhou & Turk-Browne, 2025), is that cortical entrainment supports predictive efficiency and behavioral performance, whereas representational reorganization in the hippocampus underlies the construction of subjective time. More broadly, these findings suggest that tracking the structure of input and constructing the temporal experience of that structure rely on partially separable neural processes. This aligns with previous studies showing that while cortex tracks transitional probabilities through frequency tagging, the hippocampus is insensitive to those phase-locked modulations, only showing representational change (Henin et al., 2021).

### Methodological implications and future directions

Most studies of event-based time perception rely on retrospective judgments, which conflate perception with memory, attention, and reconstruction (Block & Zakay, 1997; Grondin, 2010). By contrast, the pause-adjustment paradigm introduced here captures temporal distortion during ongoing perception, minimizing post-hoc influences. The fact that temporal warping emerged continuously while the stream unfolded provides strong evidence that event segmentation shapes the perception of time itself, not merely remembered duration. This approach therefore opens a new methodological avenue for studying the real-time construction of subjective experience.

Future work combining this paradigm with neural recordings that resolve representational changes—particularly in hippocampal and cortical circuits implicated in event segmentation—could test whether within-event compression tracks reductions in representational distance and whether boundary-related dilation corresponds to neural state resets between events. Such work would allow a direct test of the proposed link between representational geometry and subjective time.

Moreover, the events studied here are defined by transitional probabilities within a statistical learning (SL) framework, which may differ from the larger-scale, semantically-rich, or narrative-driven events (e.g., films or stories) typical of traditional event segmentation research (Zacks et al., 2007, 2011). While SL provides a controlled mechanistic baseline, it remains an open question whether the same principles govern temporal experience in more complex context e.g., those defined by high-level goals or complex narrative arcs. Extending this approach to naturalistic stimuli such as speech, music, and real-world action sequences will be essential for establishing the generality of event-based time construction.

## Conclusion

Everyday experience suggests that comprehension can slow time: the same speech stream can feel fast when unstructured and slow when parsed into meaningful units. The present experiments provide a mechanistic foothold on this intuition. When a continuous auditory stream is segmented into events through statistical learning, subjective time warps in systematic ways: boundaries expand and event interiors compress. This warping is reliable across samples of participants, persists across time scales, and is modified when events acquire semantic meaning. Together, these findings support a unified view in which subjective time is an emergent property of how experience is structured into events, linking perception, memory, and prediction within a common representational framework.

## Methods

### Experiment 1

#### Participants

A total of 41 participants took part in the study. Two participants were excluded due to low performance in the target detection task (less than 10% accuracy). The final sample was composed of 39 participants (age: 32.6 ± 8.7 years; LexTALE score: 86.3 ± 8.7, 3 native English speakers; 23 female; 36 right-handed). None of the participants self-reported hearing or visual impairments, nor had any history of drug use. All volunteers provided both oral and written informed consent before participating. All procedures were conducted in accordance with the Declaration of Helsinki. The study protocol received approval from the Ethics Council of the Max Planck Society. Participants received 7 Euros per 30 minutes of participation.

#### Experimental procedure

The experiment consisted of three phases. First, a pre-screening LexTALE task conducted online (Lemhöfer & Broersma, 2012) identified participants with high enough English proficiency. Second, an in-lab explicit statistical learning paradigm with several parts: a word-matching task, a stem-completion task, a target-detection task, followed by another round of the stem-completion task. The word-matching and the stem-completion tasks measured retention of the pseudowords in memory; the target-detection task evaluated the learning of the transitional probabilities. Third, participants completed the pause-adjustment task, which quantified time distortions resulting from statistical learning of pseudowords. The entire experiment lasted approximately 3 hours, self-paced breaks were scheduled between tasks to prevent fatigue.

The in lab part of the study was conducted in a light controlled and acoustically isolated chamber. The experiment was programmed in PsychoPy (version 2024.2.4) and data were collected on a Fujitsu CELSIUS M740B desktop computer running Windows 10 Pro. Visual stimuli were presented on a 24-inch LCD monitor with a 1920×1080 resolution and a 60 Hz refresh rate at a viewing distance of approximately 60 cm and participants’ head position was stabilized using a chinrest. Auditory stimuli were delivered binaurally through Beyer dynamics DT-770 over-ear headphones at a comfortable listening level adjusted by the participants with soundcard (RME Fireface UC) and headphone amplifier (Lakepeople G103P). Participants responded using a standard keyboard with their right hand. For the pause-adjustment task, participants controlled a visual slider using a computer mouse.

#### Stimuli

Twelve consonant-vowel syllables were generated using an online test-to-voice synthesizer (SpeechPro, Speech Technology Center, 2024) and sampled at 48000 Hz using Audacity. Syllable lengths were equated and prosody was flattened using Praat (Boersma & Weenink, 2024) to eliminate acoustic cues that could be used for parsing. The duration of each syllable was 375 ms. Individual syllables were concatenated in MATLAB (R2023b, MathWorks). Two types of sequences were created: structured streams and random streams. In the structured stream, the transitional probabilities between syllables were manipulated to create four three-syllable pseudowords: TUPIRO, GOLABU, BIDAKU and PADOTI. The transitional probabilities between syllables within a pseudoword were 1, whereas the transitional probabilities between pseudowords were 1/3, as no pseudoword could repeat back-to-back. Each pseudoword had a duration of 1,125 ms. The random stream consisted of the same 12 syllables but presented in a random order with no back-to-back syllable repeats, resulting in transitional probabilities of 1/11. Each stream lasted approximately 27 s (72 syllables, each repeated 6 times) and was presented 12 times. The timing of syllables in both streams was fixed, resulting in a syllable presentation rate of 2.67 Hz and a pseudoword presentation rate of 0.89 Hz in the structured stream (Fig. 1c).

#### LexTALE prescreening phase

The LexTALE task used in this experiment was an adapted version of the standardized task (Lemhöfer & Broersma, 2012) and was administered via Gorilla (Anwyl-Irvine et al., 2020), integrated with the local participant recruitment system. On each trial, a word or non-word was presented on the screen for 5 s and participants were asked to decide whether it is an existing English word. There were a total of 40 words and 20 non-words (60 trials), and the words and their order of appearance on each trial strictly followed the instructions of the standardized LexTALE task (Lemhöfer & Broersma, 2012). As this was an online task, we took extra precautions to identify and remove trials and participants based on their performance. Specifically, all responses exceeding 5s on a trial were considered incorrect. Additionally, all words were displayed as images on the screen to avoid copying and translation. We used the mean accuracy of words and non-words to detect English proficiency (Lemhöfer & Broersma, 2012), and only participants with a score of 70 were invited to the main in-lab experiment.

#### Explicit statistical learning phase

Subjects were pre-trained on the four pseudowords before exposure to the structured stream in order to facilitate parsing and maximize potential effects on time perception. In the word-matching task, participants listened to pseudowords and chose the word they had heard from four written options displayed horizontally on the screen (Fig. 1a). They were encouraged to memorize the spelling and pronunciation of the pseudowords because their memory would be tested after the study. The word-matching task had a total of 40 trials, 10 for each pseudoword. Participants completed a stem-completion task, in which they were presented with one syllable of a pseudoword and had to type the corresponding pseudoword. There were 12 trials, with each syllable serving as the stem once (Fig. 1b). Participants who could complete fewer than 8 pseudowords repeated the word-matching task and tried again. Participants who failed the stem-completion task three times were excluded from the study due to low memory performance (16 out of 57 were excluded, N=41 completed all tasks).

After participants explicitly learned the pseudowords, they were exposed to the structured and random streams. To ensure attention to the streams, and to evaluate the learning benefits in the explicit statistical learning condition, participants completed a target-detection task (Batterink, Reber, & Paller, 2015).

Before each stream, they were presented with a syllable and asked to detect it in the upcoming stream by pressing the spacebar as quickly as possible (Fig. 1d). Response time as well as accuracy were emphasized. The structured and random streams were presented in separate blocks, with the block order counterbalanced across participants. Each block consisted of 12 trials, in which one syllable served as the target, for a total of 24 trials. Retention of the pseudowords was measured once more (stem-completion task) at the end of this phase without any performance requirement.

#### Pause-adjustment phase

To quantify time distortions in a continuous auditory stream as a function of learning, we developed an online pause-adjustment task (Fig. 1e). We hypothesized that time perception would be differentially distorted for pauses occurring within pseudowords versus between pseudowords, an effect expected to arise only for the structured stream. Such distortions could manifest as temporal compression or expansion, potentially explaining the subjective impression that the stream slowed down or adopted a different cadence.

To measure these distortions, participants were instructed to adjust the durations of pauses in the stream until the rhythm sounded monotonous or temporally regular across all syllables. This procedure allowed us to estimate each participant’s point of subjective equality (PSE) for pause duration, providing a behavioral measure of time distortion. Across trials, either the pauses within pseudowords were held constant while participants adjusted the pauses between pseudowords, or vice versa.

During each trial, participants used a slider displayed on the screen to adjust pause duration continuously while the stream was played without interruption. The slider provided real-time auditory feedback: changes to the slider instantaneously altered the duration of the relevant pauses, allowing participants to directly hear the effect of their adjustments, analogous to adjusting volume while listening to music. Participants could move the slider as many times as needed and terminate the trial by pressing the spacebar once they perceived all pauses as having equal duration. This continuous playback ensured that performance relied on online time perception rather than memory.

The structured session comprised two conditions: a *within-word* condition and a *between-word* condition. In both conditions, a structured stream was presented on each trial, initially configured with pauses between words that were either noticeably longer or shorter than pauses within words. In the *within-word* condition, pauses within pseudowords served as the fixed reference and could not be adjusted; participants were instructed to adjust the pauses between pseudowords until they matched the perceived duration of the within-word pauses. This condition was termed *within-word* because the adjustment readout reflects perceived within-word timing. Conversely, in the *between-word* condition, pauses between pseudowords were held constant as the reference and participants adjusted the pauses within pseudowords to match them. Participants were blinded to the condition labels to minimize bias. By comparing adjustments across these two conditions, we assessed how event segmentation influenced perceived duration for within-and between-word intervals.

To control for potential biases arising from the experimental structure itself, we included a random session using random syllable streams lacking transitional structure. To match the structured session as closely as possible, we arbitrarily grouped every three consecutive syllables into a “pseudoword” and applied the same pause adjustment procedures, now dummy coded. The two conditions in this session were termed *fake within-word* and *fake between-word* conditions, in analogy to the structured conditions.

To examine the generalizability of time distortions, we manipulated the reference pause duration across four levels (35, 55, 75, 95 ms). For each session, condition, and reference level, participants completed 16 trials, yielding a total of 128 trials per session.

For the pause-adjustment task, we constructed 60 structured streams and 60 random streams. Each stream consisted of 216 syllables (corresponding to 72 pseudowords in the structured streams). Unlike the stimuli used in the target-detection task, these streams were not pre-generated. Instead, they were constructed dynamically in real time during the experiment. Pauses between syllables were inserted online as syllables were played, allowing their duration to be flexibly adjusted according to the experimental condition and participant input.

## Data analysis

### Explicit memory: word-matching and stem-completion tasks

To assess explicit learning and retention of the pseudowords, we analyzed performance on the stem-completion tasks at two time points: immediately after the initial word-matching training and again after completion of the exposure to the structured and random streams. Mean accuracy was computed for each participant at each time point. Paired-sample t-tests were used to compare accuracy between the two tests, providing a measure of memory persistence across the experiment. Unless otherwise explicitly stated, all statistical tests were two-sided. For paired-samples t-tests, Cohen’s *d_z_* is reported, which was calculated by dividing the mean difference by the standard deviation of the difference scores, appropriate for within-subject designs.

### Online statistical learning: target-detection tasks

Online sensitivity to the statistical structure of the auditory streams was assessed using response times (RTs) from the target-detection task. Only responses 100-1500 ms after the target syllable onset were included; responses outside this window were classified as false alarms and excluded from RT analyses. For each participant, mean RTs were computed per syllable position within pseudowords (first, second, third syllable). These values were entered into a repeated-measures ANOVA with syllable position as a within-subject factor separately for the structured and random streams. Significant main effects were followed by post-hoc paired t-tests to compare detection speed between specific syllable positions. The assumption of sphericity was evaluated using Mauchly’s test, degrees of freedom were adjusted using the Greenhouse-Geisser correction if violated.The magnitude of statistical learning for each participant was quantified using a normalized RT-based prediction index. This index was defined as the difference between RTs to first syllables and the mean RTs to second and third syllables, normalized by their sum. Larger values indicate stronger RT facilitation for predictable syllables.

To evaluate whether sequential RT facilitation was specific to the learned event structure, these mean RTs were also entered into a two-way repeated-measures ANOVA with stream type (structured vs. random) and syllable position (first, second, third) as within-subject factors. Moreover, the RT-based prediction indices were directly compared between the structured and random streams using a paired-samples t-test. For repeated-measures ANOVAs, partial η^2^ is reported. For paired-samples t-tests, Cohen’s *d_z_*is reported. *BF*_10_ was reported for non-significant comparisons to explicitly quantify the evidence in favor of the null hypothesis, with Default Cauchy prior width of *r* = 0.707.

### Subjective time perception: pause-adjustment task

Subjective time distortions were quantified using pause adjustments. For each trial, the adjusted pause duration was recorded. To isolate perceived distortions relative to the physical stimulus, we computed relative response pauses by subtracting the reference pause duration from the adjusted pause duration.

We employed linear mixed-effects models (LMMs) in R (version 4.4.1) using the lmerTest package (v3.1.3, Kuznetsova et al., 2017) to examine how statistical learning influenced subjective time perception while accounting for inter-individual variability. Degrees of freedom and *p* values for the LMMs were estimated using Satterthwaite’s approximation. In the initial model, relative response pauses were pooled across reference levels to assess the overall distortion effect. Fixed effects included session (structured vs. random), condition (within-word vs. between-word), and their interaction. Participants were included as a random intercept. No demographic or cognitive covariates were included in the statistical models.

Significant interactions were followed by Holm-corrected simple effects analyses.

To evaluate whether time distortions generalized across different temporal scales, a second LMM was constructed using absolute adjusted pause durations as the dependent variable. This model included reference level (35, 55, 75, 95 ms) as an additional fixed effect and tested for a three-way interaction between session, condition, and reference level.

As a complementary analysis, we computed ratios of adjusted pauses in the between-word condition relative to the within-word condition for each reference level. One-sample t-tests with false discovery rate (FDR) correction (Benjamini & Hochberg, 1995) were used to test whether these ratios were significantly greater than 1, indicating that pauses between pseudowords were perceived as longer than pauses within pseudowords.

### Relationship between statistical learning and time distortion

To examine whether individual differences in statistical learning predicted the magnitude of subjective time distortion, Pearson correlation analyses were conducted between the normalized RT-based prediction index derived from the target-detection task and a time-distortion index derived from the pause-adjustment task. The time-distortion index was computed by baseline-correcting adjusted pauses with their corresponding pauses from the random condition, rescaling these values to a range of 0-1 across participants and conditions, and calculating a normalized contrast of between-word vs. within-word conditions, averaged across reference levels. In addition to frequentist correlations, Bayesian Pearson correlations were performed to quantify evidence for the absence of a relationship between statistical learning and time distortion. Bayes factors (*BF*_10_) with default Cauchy prior width of *r* = 0.707.

### Experiment 2

#### Participants

A total of 37 participants (mean age = 29.7 ± 6.5 years; LexTALE score = 84.5 ± 8.4, 8 native English speakers; 15 female; 36 right-handed) completed the experiment, using the same recruitment criteria as in Experiment 1. None of the participants self-reported hearing or visual impairments, nor had any history of drug use. All participants had no history of neurological disorders. All volunteers provided oral and written informed consent prior to participation. The study was conducted in accordance with the Declaration of Helsinki and approved by the Ethics Committee of the Faculty of Medicine at Goethe University. Participants received monetary compensation of 10 Euros per 30 minutes of participation.

All participants met the behavioral inclusion criteria, as applied in Experiment 1. Magnetoencephalographic data were collected in a 275-channel CTF MEG systems (VSM MedTech Omega 2005) for all participants in a magnetic isolated faraday cage (those data will be reported in other publications). In addition, pupil diameter was recorded during the explicit statistical learning phase to assess whether auditory regularities elicited frequency-specific oscillations in pupil size, used here as a physiological index of event segmentation. We further examined whether the strength of pupil entrainment was related to individual differences in event segmentation and to the magnitude of temporal distortion observed in the pause-adjustment task.

Pupil data were collected during the target-detection task using an eye-tracker. Gaze position and pupil size were recorded continuously from the right eye using a video-based system (EyeLink Portable Duo, SR Research) at a sampling rate of 1000 Hz. A nine-point calibration procedure was performed before each block. To ensure data quality for the pupil entrainment analyses, we calculated the percentage of pupil data loss for each block. Blocks were classified as low quality if more than 25% of pupil samples were lost due to blinks or noise. Participants with fewer than six usable blocks were excluded from pupil analyses. This criterion ensured sufficient valid data across both experimental conditions. Fourteen participants were excluded on this basis. The final sample for pupil analyses therefore consisted of 23 participants (mean age = 28.7 ± 5.7 years; 10 female; LexTALE score = 85.4 ± 8.3, 3 native English speakers; all right-handed). The total duration of the experimental session was approximately 3 hours.

#### Stimuli and experimental procedure

Experiment 2 followed the same general procedure as Experiment 1, with the following modifications. First, eye-tracking and pupil data were collected to quantify entrainment to the structure of the auditory streams and to obtain an individualized physiological measure of learning. Second, to enable frequency-domain analyses of the pupil signal, the duration of the auditory streams was increased to 81 s, corresponding to 216 syllables and 72 trisyllabic pseudowords in the structured streams. Third, MEG data were collected during this experiment but will be reported in a separate publication.

The experiment was programmed in Psychopy (version 2024.2.4), and data were collected on a costumed Krotus desktop computer (Mainboard: ASUS ProArt B760 CREATOR; CPU: Intel Core i9-14900K) running Windows 11. Visual stimuli were presented using a VPixx ProPixx laser projector (1920 × 1080 resolution, 60 Hz refresh rate), located outside the MEG booth and viewed via a mirror system (dimensions: 30.7 x 23cm) with a viewing distance of approximately 50 cm. Auditory stimuli were delivered monaurally through Eartone 3A air-conduction transducers, driven by an amplifier (Lake People Phone-Amp G109 S/P) and an RME Fireface UCX soundcard. Behavioral responses were collected using a non-magnetic, fiber-optic 8-channel response pad designed for MEG environments.

Participants used the upper left and right buttons to control the slider on the screen during pause adjustment task.

Participants first learned the pseudowords via the matching task, followed by the stem-completion task. Then they completed the target detection task as well as the pause adjustment task. Unlike Experiment 1, where these tasks were completed in separate blocks, the current study interleaved target-detection and pause-adjustment tasks to minimize interference between structured and random streams.

In total, each stream type (structured and random) comprised 12 target-detection trials and 96 pause-adjustment trials. The latter included all combinations of a subset of the reference pause durations from Experiment 1 (55, 75 ms) and adjustment conditions (within-word vs. between-word). With each combination presented twice, this yielded 96 adjustment trials per condition. Within each block, participants followed a repeating trial combination: one target-detection trial followed by 8 pause-adjustment trials, and this trial combination was repeated 3 times in the same block. Each participant completed four blocks per stream type (structured and random), with the order of stream type counterbalanced across participants. The same stem-completion task from Experiment 1 was administered both before the first block and after the fourth block to assess memory persistence.

## Data Analysis

Unless otherwise noted below, Experiment 2 used the same analysis procedures as Experiment 1.

### Explicit memory: word-matching and stem-completion tasks

Besides compresence between the memory performance for the initial test and the middle test, we also performed mixed-design ANOVA with Timepoint (initial test vs. middle test) as the within-subject factor and Experiment (Experiment 1 vs. Experiment 2) as the between-subject factor to evaluate whether memory performance and retention differed across experimental cohorts. For the mixed-design ANOVA, η ^2^ was reported as the measure of effect size. For any between-subject comparisons across the two experiments, Cohen’s *d* was reported, calculated using the pooled standard deviation of the two groups.

### Online statistical learning: target-detection tasks

The normalized RT-based prediction indices derived from the structured streams were directly compared between Experiment 1 and Experiment 2 using an independent-samples t-test.

### Pupil preprocessing

Raw pupil diameter was first extracted from the right eye at the native sampling rate of the eye tracker. All preprocessing steps and analyses were conducted in MATLAB (R2023b, MathWorks). For each participant, blinks and signal dropouts were identified using the noise-based blink detection algorithm (Hershman et al., 2018). Linear interpolation was performed within a window surrounding each blink (100 ms before blink onset and 200 ms after blink offset) to ensure smooth transitions. Because artifacts may arise from transient noise unrelated to blinks, we further detected outliers using a sliding-window outlier detection procedure. For each sample, the local mean and standard deviation were calculated within a 5-s centered sliding window. Samples deviating by more than 3 times standard deviation from the local mean were identified as outliers. These segments were again linearly interpolated.

Following interpolation, the pupil signal was low-pass filtered at 5 Hz using a Kaiser-sinc windowed FIR filter (filter order = 1812), matching the filtering strategy used in prior pupillometry work (Schwiedrzik & Sudmann, 2020). To remove slow drifts in pupil diameter, we additionally applied a zero-phase 0.1-Hz high-pass Butterworth filter, which ensured that long-term fluctuations unrelated to task structure did not bias subsequent analyses.

The cleaned continuous pupil signal was segmented into separate blocks for each stream presented in the target-detection task. The resulting blocks were detrended, demeaned, and z-scored relative to the 1.5s baseline period before the start of the first syllable in the stream. Each block was evaluated for data quality by computing the percentage of interpolated data. Blocks containing > 25% interpolated data were excluded from further analyses.

### Pupil spectral power analysis

To assess whether pupil dynamics entrained to the temporal structure of the auditory streams, spectral power analyses were conducted on block-epoched pupil data. Power spectral density (PSD) was estimated using Welch’s method with a Hanning window (length = 16,875 samples, 50% overlap, 16,875 DFT points), yielding sufficient frequency resolution to capture syllable-level (2.667 Hz) and word-level (0.889 Hz) frequencies (Welch, 1967).

Because pupil spectra exhibit substantial aperiodic (1/f-like) structure that can obscure narrowband peaks, we applied Fitting Oscillations and One-Over-F (FOOOF) to separate periodic and aperiodic components (Donoghue et al., 2020). FOOOF was fit over the range 0.1–4 Hz. The fitted aperiodic component was subtracted from the original spectrum, yielding an aperiodic-corrected PSD. The resulting corrected spectra isolated narrowband periodic components and improved sensitivity to subtle entrainment effects. For each participant, FOOOF-corrected power spectra were averaged across retained blocks separately for structured and random conditions.

Statistical analyses focused on power at two theoretically expected frequencies, at the rate of the words (0.889 Hz) and the syllables (2.667 Hz). To determine whether the spectra exhibited reliable peaks at these frequencies, power was compared against a local baseline defined as the mean of the two adjacent frequency bins on either side, using paired t-tests. Relative peak power was computed by subtracting this local baseline and used in subsequent analyses. Cohen’s *d_z_* and *BF*_10_ for non-significant results were reported.

To assess condition-and frequency-specific effects, relative peak power was submitted to a two-way repeated-measures ANOVA with condition (structured, random) and frequency (word, syllable) as within-subject factors. Significant interactions were followed by planned simple-effects analyses. For repeated-measures ANOVAs, partial η^2^ was reported. For paired-samples t-tests, Cohen’s *d_z_* was reported.

*BF*_10_ was reported for non-significant comparisons.

### Pupil phase coherence analysis

We also computed inter-trial phase coherence (ITPC) to assess consistent phase-locking of pupil dynamics to the syllable and word frequencies. To this end, we segmented the preprocessed continuous pupil recordings from each block into non-overlapping pseudotrials of 15.75 s (corresponding to 42 syllables or 14 pseudowords), aligned to the onset of the first syllable in a pseudoword. This segmentation ensured sufficient frequency resolution to distinguish ITPC at the syllable and word levels. To ensure data quality, only trials with < 25% interpolated samples were included in the analysis.

We next computed a trial-by-trial Fourier transform of the segmented pupil pseudotrials using single discrete prolate spheroidal sequence (DPSS) as a taper, applying the same frequency resolution as in the power spectrum analyses to ensure direct comparability across measures. This analysis was implemented using the FieldTrip toolbox (version 20250402; Oostenveld et al., 2011). ITPC was then calculated for each participant and condition, with the same method as used in previous study (Schwiedrzik & Sudmann, 2020). Elevated ITPC at the word (0.889 Hz) or syllable (2.667 Hz) frequencies indicates phase-locked entrainment of pupil dynamics.

To determine whether ITPC at the target frequencies exceeded chance levels, we performed a phase-randomization permutation test. For each participant, the complex Fourier coefficients at the frequency of interest were rotated by random phase offsets and ITPC was recomputed 1,000 times to obtain a participant-specific null distribution (preserving spectral amplitude but eliminating cross-trial phase alignment). The null ITPC for each frequency and condition was defined as the mean of this distribution. Observed ITPC values were then compared against the null values using two-tailed paired t-tests. Cohen’s *d_z_* and *BF*_10_ for non-significant results were reported.

To assess how ITPC differed across conditions and frequencies, ITPC values were submitted to a two-way repeated-measures ANOVA with condition (structured, random) and frequency (word, syllable) as within-subject factors. Significant interactions were followed up using planned simple-effects contrasts comparing structured and random conditions at each frequency. For repeated-measures ANOVAs, partial η^2^ was reported. For paired-samples t-tests, Cohen’s *d_z_* was reported. *BF*_10_ was reported for non-significant comparisons.

### Behavior analysis and pupil-task correlations

To examine whether physiological entrainment reflected individual sensitivity to statistical structure, we computed Pearson correlations between word-level ITPC in the structured condition and the normalized RT-based prediction index derived from the target-detection task.

To test whether physiological entrainment predicted subjective time distortion, we correlated word-level ITPC with the magnitude of the pause-distortion effect. For each participant, the distortion effect was quantified as the difference between adjusted pause durations in the between-word and within-word conditions, pooled across reference levels. Pearson correlations were used to assess these relationships.

*BF*_10_ was reported for non-significant result.

### Experiment 3

Experiment 3 introduced semantic associations for each pseudoword to examine whether the addition of meaning enhances event segmentation and modulates its influence on subjective time perception.

## Method

### Participants

To control for potential effects of native language on semantic associations, we recruited native German speakers with at least B1-level English proficiency, as assessed by the LexTALE task. A total of 53 participants completed the experiment. None of the participants self-reported hearing or visual impairments, nor had any history of drug use. All volunteers provided oral and written informed consent prior to participation. The study was conducted in accordance with the Declaration of Helsinki and approved by the Ethics Council of the Max Planck Society. Participants received 7 Euros per 30 minutes of participation.

All participants met the behavioral inclusion criteria. Fourteen participants were excluded from the pupil data analyses using the same criteria applied in Experiment 2. The final sample included in the eye-tracking analyses therefore consisted of 39 participants (mean age = 32.2 ± 9.2 years; LexTALE score = 85.8 ± 9.3; 23 female; all right-handed).

### Stimuli

Experiment 3 used the same set of consonant-vowel syllables, pseudowords, and auditory stream configurations as in Experiment 2. For the semantic association task, four trisyllabic German nouns were selected to serve as meanings for the four pseudowords. Word selection was based on lexical frequency measures extracted from the SUBTLEX-DE corpus to ensure comparability across items (Brysbaert et al., 2011). In addition, semantic similarity and dissimilarity were quantified using CLIP-based embeddings to ensure that the selected meanings spanned distinct conceptual categories (Radford et al., 2021). The final set of meanings consisted of *basketball*, *cassette*, *hamburger*, and *squirrel*. The images used to depict these meanings were drawn from the THINGS database (Hebart et al., 2019).

The in lab part of the study was conducted in a light controlled and acoustically isolated chamber. The experiment was programmed in PsychoPy (version 2024.2.4) and data were collected on a Fujitsu ESPRIMO P958 desktop computer running Windows 10 Enterprise. Visual stimuli were presented on a 24-inch LCD monitor with a 1920×1080 resolution and a 60 Hz refresh rate at a viewing distance of approximately 60 cm and participants’ head position was stabilized using a chinrest. Auditory stimuli were delivered binaurally through Beyer dynamics DT-770 Pro over-ear headphones at a comfortable listening level adjusted by the participants with soundcard (RME Fireface UC) and headphone amplifier (Lakepeople G103P). Participants responded using a standard keyboard with their right hand. For the pause-adjustment task, participants controlled a visual slider using a computer mouse.

Pupil size was continuously recorded from the right eye during the target-detection and pause-adjustment tasks using a video-based eye-tracker (EyeLink 1000 Plus, SR Research) at a sampling rate of 1000 Hz. A nine-point calibration was performed before the start of each block. Trigger signals were used to synchronize stimulus events with physiological recordings.

### Experimental procedure and data analysis

As in Experiments 1 and 2, participants first completed a pseudoword pre-training phase using the word-matching task prior to the structured and random blocks. In Experiment 3, following successful learning of the pseudowords, participants additionally completed a semantic association learning task. For each participant, the four meanings were randomly assigned to the four pseudowords (Fig. 3a).

In each semantic learning trial, participants heard a pseudoword while viewing an image depicting its assigned meaning and then selected the correct pseudoword from a set of four alternatives. Participants were instructed to memorize both the phonological form and the associated meaning of each pseudoword. Following this matching phase, participants were shown an image and asked to reconstruct the corresponding pseudoword by selecting its constituent syllables in the correct order from a randomized list. Accuracy feedback was provided after each trial, and learning cycles continued until participants achieved perfect recall of all pseudoword–meaning pairings. This semantic memory task was repeated after completion of four statistical learning and pause-adjustment task blocks to assess retention.

Participants completed three blocks per stream type (structured and random), with stream order counterbalanced across participants (three structured blocks followed by three random blocks, or vice versa). Within each block, participants first completed two trials of the target-detection task. They then performed the pause adjustment task with all combinations of pause reference duration (35, 55, 75, 95 ms) and adjustment condition (within-word vs. between-word), with each combination presented three times. These paired target-detection and pause-adjustment trials were repeated twice per block. This resulted in a total of 12 target-detection trials and 144 pause-adjustment trials per participant, evenly distributed across structured and random conditions. The total duration of the experimental session was approximately 3.5 hours.

Behavioral and pupillometry data from Experiment 3 were preprocessed and analyzed using the same procedures as in Experiment 2.

Besides the standard procedures, we performed a series of comparisons between the cohort in Experiment 3 (N = 53) and the pooled dataset from Experiments 1 and 2 (N = 76).

Differences in memory performance and the magnitude of statistical learning (normalized RT-based prediction indices) between these cohorts were evaluated using the identical mixed-design ANOVA and independent-samples t-test procedures described in Experiment 2. As previously specified, η_G_ ^2^and Cohen’s *d* are reported as effect sizes for these analyses, respectively.

Additionally, to determine if semantic enrichment modulated the overall extent of temporal warping, independent sample t-test was conducted to compare the normalized pause-distortion indices between the Experiment 3 cohort and the pooled Experiments 1 and 2 cohort. Consistent with previous between-subject analyses, a *BF*_10_ was reported using a default Cauchy prior width of r = 0.707.

## Acknowledgements

We thank the members of the NCC and the PB lab for their feedback. C.M. Schwiedrzik for insightful discussions and feedback throughout the different phases of this study, as well as for sharing the code for pupil analyses. We thank T. Öztürk, J. Taube, N. Kabirmokhtarfar, N. Arabshahi, M. Eisenberg, N. H. T. Tran, M. Schütz and M. Wittstock for assistance with piloting and data collection. This work was supported by Bial Foundation (No. 268/22 to L.M and D.T), CIFAR catalyst grant (CF-0434-CP24-071 to L.M and N.B.T.-B.), Marie-Curie Individual Fellowship (No. 101023805 to D.T.), the China Scholarship Council (No. 202306750009 to Q.Z.) and the Max Planck Society.

## Authors Contribution

Conceptualization: D.T., N.B.T.-B., L.M. Methodology: Q.Z., D.T., N.B.T.-B., L.M. Software: Q.Z. Validation: Q.Z., D.T., L.M. Formal analysis: Q.Z. Investigation: Q.Z. Resources: L.M. Data curation: Q.Z. Writing-original draft: Q.Z., L.M. Writing-review and editing: Q.Z., D.T., N.B.T.-B., L.M. Visualization: Q.Z., L.M. Supervision: D.T., L.M. Project administration: L.M. Funding acquisition: D.T., N.B.T.-B., L.M. All authors reviewed and approved the final version of the manuscript.

## Data availability

The processed data are deposited at Zenodo, which has not yet been publicly released.

## Code availability

The analysis scripts and figure-generation code used in this study are deposited at Zenodo, which has not yet been publicly released.

## Competing Interests

The authors declare no competing interests.

## Supplementary Information

**Supplementary Fig. 1.**
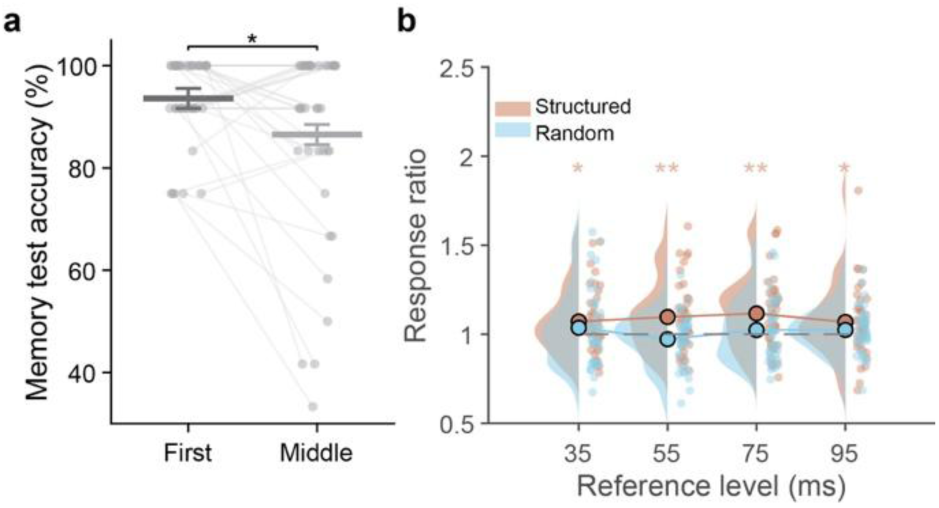
Robust temporal warping of subjective time in Experiment 1. **a**, Explicit memory for the four trisyllabic pseudowords assessed immediately after learning (First) and midway through the experiment (Middle). Although accuracy declined significantly, performance remained high at both time points. **b**, Ratios of adjusted pauses (between-word / within-word). Ratios were significantly greater than 1 at all reference levels in the structured stream, indicating reliable boundary-related expansion across time scales. Individual scatter points and violin distributions represent raw data to preserve the original scale. Error bars in panel (**a**) denote Morey-corrected SEM. *P < 0.05; **P < 0.01. To assess robustness across time scales, we computed the ratio of adjusted between-word to within-word pauses separately for each reference duration. Ratios greater than 1 indicate longer perceived pauses at event boundaries. All reference levels showed ratios significantly above 1 after FDR correction (35 ms: *t*(38) = 2.44, *p*_FDR_ = 0.026; 55 ms: *t*(38) = 3.06, *p*_FDR_ = 0.008; 75 ms: *t*(38) = 3.55, *p*_FDR_ = 0.004; 95 ms: *t*(38) = 2.12, *p*_FDR_ = 0.041), confirming that the temporal distortion generalized across pause durations (Supplementary Fig. 1b).

**Supplementary Fig. 2.**
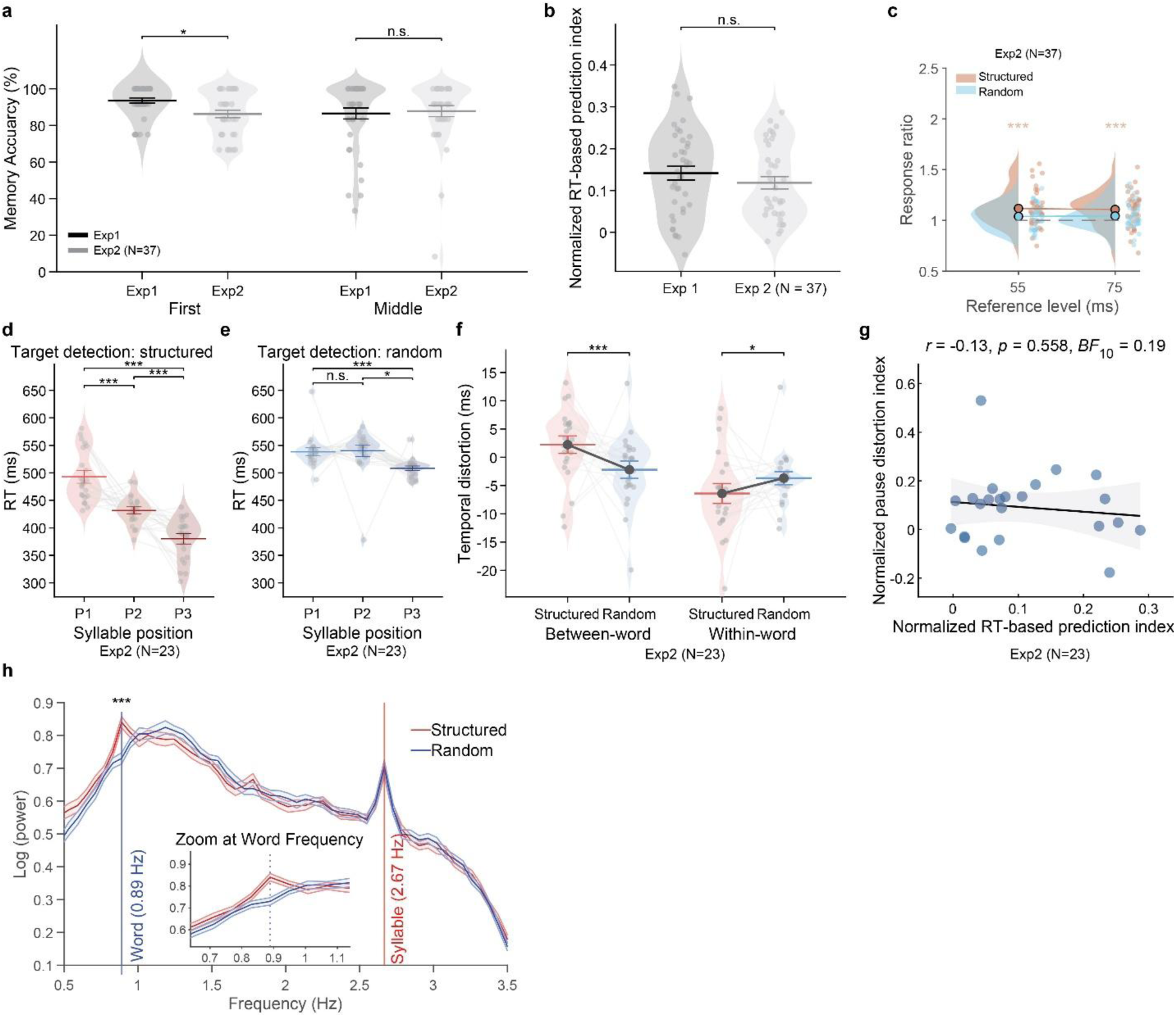
Behavioral replication, combined analyses, and pupillary power spectra in Experiment 2. **a**, Memory accuracy for the trisyllabic pseudowords at initial training (First) and midway through the experiment (Middle) for Experiment 1 (N = 39) and Experiment 2 (N = 37). Mixed ANOVA showed marginally significant interaction between experiment and test time (initial vs. midway; *F*(1,74) = 3.956, *p* = 0.050, η_G_² = 0.021). Experiment 1 showed higher initial memory accuracy than Experiment 2 (*t*(141) = 2.115, *p* = 0.036, Cohen’s *d* = 0.69, CI: [0.48, 14.18]), but no difference was found at the midway test (*t*(141) =-0.375, *p* = 0.708, Cohen’s *d* = 0.07, CI: [-8.15, 5.55], *BF*_10_ = 0.247). In another direction, within Experiment 1, responses significantly decreased from first-test to middle-test (*t*(74) =-2.330, *p* = 0.023, Cohen’s *d_z_* = 0.41, CI: [-13.08,-1.02]). In contrast, Experiment 2 showed no significant change over time (*t*(74) = 0.507, *p* = 0.613, Cohen’s *d_z_* = 0.08, CI: [-4.62, 7.77], *BF*_10_ = 0.196). **b**, The normalized RT-based prediction index showed no significant difference between Experiment 1 and Experiment 2, indicating consistent learning across cohorts. **c**, Ratios of adjusted pauses (between-word / within-word) were significantly greater than 1 for both reference levels in the structured session, confirming a stable distortion across temporal scales (N = 37). **d–h**, Behavioral and physiological results for the eye-tracking subsample (N = 23). **d**, Target-detection RTs in the subsample showed significant modulation by syllable position in structured streams. **e**, Such sequential modulation was absent in the random streams. **f**, Differences between structured and random sessions in different conditions. In the structured session, pauses at event boundaries (between-words) were perceived as longer (temporal expansion), whereas pauses within events (within-words) were perceived as shorter (temporal contraction). **g**, No significant correlation was observed between the implicit prediction index and the magnitude of pause distortion. **h**, FOOOF-corrected power spectra of pupil diameter revealed significant entrainment at both the word (0.889 Hz) and syllable (2.667 Hz) frequencies in structured streams. In random streams, a significant peak emerged only at the syllable frequency. Individual scatter points and violin distributions represent raw data to preserve the original scale for between-subject analyses (panel a, b), and represent within-subject normalized values computed via the Cousineau-Morey method (panel d-f). Error bars in panels (a, b) represent SEM, and represent Morey-corrected SEM in panels (d-f, h). Shaded areas in the correlation panel (**f**) represent the 95% Confidence Interval (CI). *P < 0.05; **P < 0.01; ***P < 0.001; N.S., not significant. Before examining pupil dynamics, we verified that the Experiment 2 cohort with eye-tracking exhibited the same behavioral signatures of learning as the full cohort. The cohort retained a robust modulation of RTs by syllable position in structured streams (Greenhouse-Geisser corrected *F*(1.48, 32.56) = 34.78, *p* < 0.001, partial η^2^ = 0.61; Supplementary Fig. 2d), controlling by random streams (Supplementary Fig. 2e) further confirming successful statistical learning. Furthermore, the event-driven distortion of subjective time replicated in this group. An LMM analysis confirmed a significant interaction between session (structured vs. random) and condition (within-words vs. between-words) (*β* =-0.007, *t*(4390) =-3.932, *p* < 0.001, CI: [-0.011,-0.004]), driven by the expansion of pauses between words (Estimate = 0.004, *t*(4390) = 3.454, *p* < 0.001, CI: [0.002, 0.007]) and compression of pauses within words (Estimate = - 0.003, *t*(4390) =-2.106, *p* = 0.035, CI: [-0.005,-0.001]) in the structured session relative to the random session (Supplementary Fig. 2f). Finally, no significant correlation was observed between the RT-based prediction index and the magnitude of the pause-distortion effect (*r* =-0.129, *p* = 0.558, CI: [-0.514, 0.299], *BF*_10_ = 0.19 < 1/3, Supplementary Fig. 2g). These findings confirm that the eye-tracking cohort in Experiment 2 remains behaviorally representative, providing a valid basis for investigating the physiological correlates of these effects. To investigate whether pupil size entrained to the statistical structure of auditory streams, we also analyzed the spectral power of pupil diameter signals during the target-detection task. After removing aperiodic components using FOOOF, corrected power spectra revealed peaks at both the syllable (2.667 Hz) and word (0.889 Hz) frequencies in the structured streams. Paired *t*-tests comparing the power at each frequency to the surrounding baseline bins showed significant entrainment at the syllable frequency (*t*(22) = 4.763, *p* < 0.001, CI: [0.08, 0.20], Cohen’s *d_z_* = 0.88) and the word frequency (*t*(22) = 5.413, *p* < 0.001, CI: [0.05, 0.11], Cohen’s *d_z_* = 0.38). In contrast, for the random streams, a significant peak was observed only at the syllable frequency (*t*(22) = 5.842, *p* < 0.001, CI: [0.10, 0.21], Cohen’s *d_z_* = 1.03), while power at the word frequency did not differ from baseline (*t*(22) =-1.426, *p* > 0.05, CI: [-0.03, 0.01], Cohen’s *d_z_* = 0.08, *BF*_10_ = 0.556). A two-way repeated-measures ANOVA with frequency (syllable, word) and stream type (structured, random) revealed a significant interaction (*F*(1, 22) = 23.464, *p* < 0.001, partial η^2^ = 0.52). A post-hoc test showed significantly higher relative power for structured streams at the word frequency (*t*(22)=5.788, *p* < 0.001, CI:[0.06, 0.13], Cohen’s *d_z_* = 1.23), but not the syllable frequency (*t*(22)=-0.963. *p* = 0.346, CI: [-0.05, 0.02], Cohen’s *d_z_* = 0.21, *BF*_10_ = 0.331 < 1/3), confirming that frequency tagging at the word level was specific to structured input (Supplementary Fig. 2h).

**Supplementary Fig. 3.**
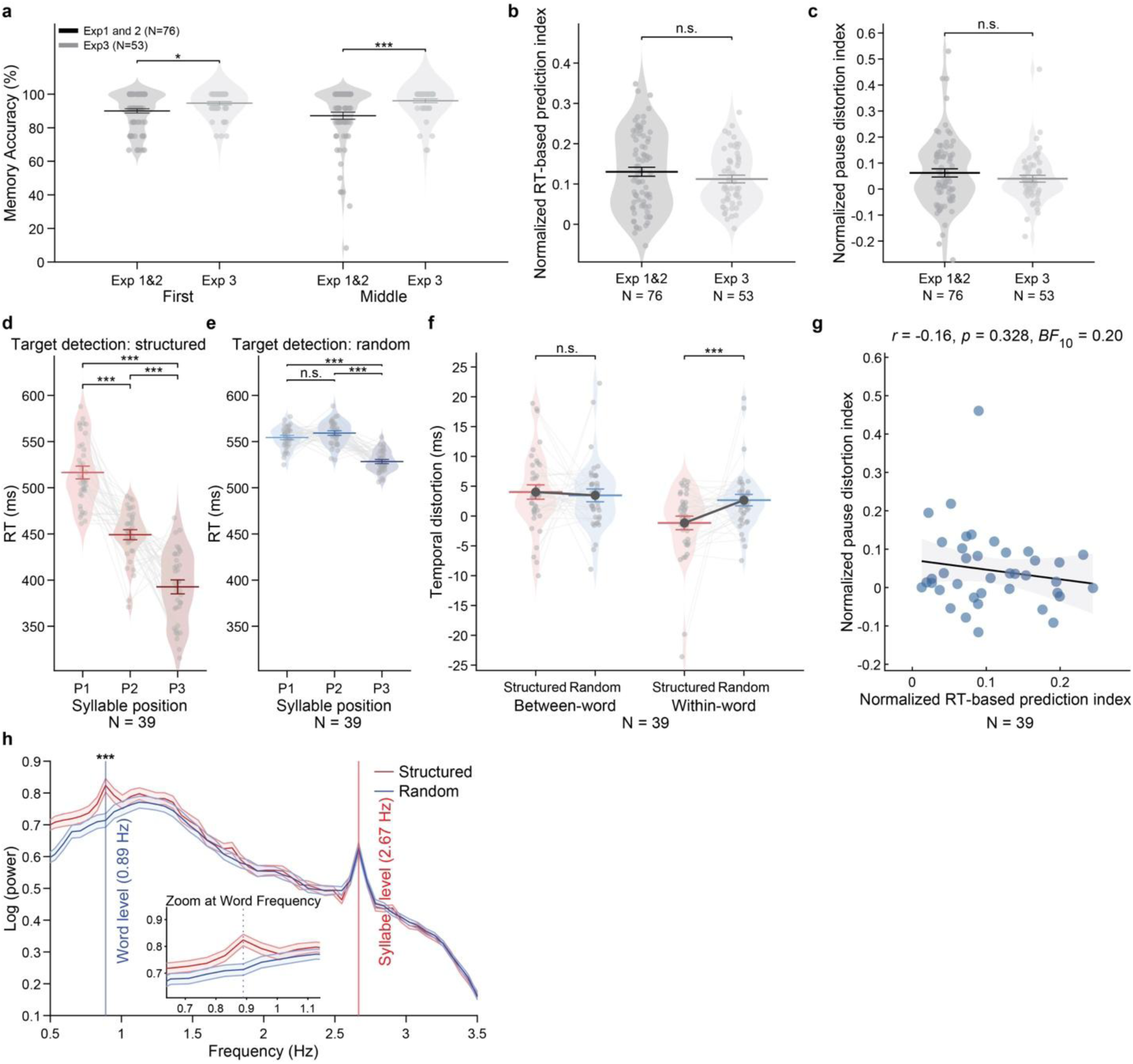
Enhanced memory stability and shifted temporal distortion under semantic enrichment. **a**, Accuracy for the trisyllabic pseudowords at the midway test was significantly higher in Experiment 3 (N = 53) than in the combined sample of Experiments 1 and 2 combined (N = 76). **b**, The normalized RT-based prediction index showed no significant difference between Experiment 3 and the combined results of Experiments 1 and 2, indicating that semantic enrichment did not alter the basic extraction of regularities. **c**, The magnitude of the temporal distortion effect (normalized difference between between-word and within-word pauses) did not differ significantly between Experiment 3 and the combined sample of Experiment 1 and 2. **d-h**, Behavioral and physiological results for the cohort with eye-tracking in Experiment 3 (N = 39). **d**, RTs showed a robust modulation by syllable position in structured streams, confirming successful statistical learning in this cohort. **e**, Such modulation was absent in the random streams. **f**, Differences between structured and random sessions show that semantic meaning eliminates boundary expansion. **g**, No significant correlation was observed between the behavioral RT-based prediction index and the magnitude of subjective pause distortion, replicating the dissociation seen in Experiments 1 and 2. **h**, FOOOF-corrected power spectra of pupil diameter revealed clear peaks at both word (0.89 Hz) and syllable (2.67 Hz) frequencies in structured streams, while random streams exhibited entrainment only at the syllable frequency. Individual scatter points and violin distributions represent raw data to preserve the original scale for between-subject analyses (panel a-c), and represent within-subject normalized values computed via the Cousineau-Morey method (panel d-f). Error bars in panels (a-c) represent SEM, and represent Morey-corrected SEM in panels (d-f, h). Shaded areas in the correlation panel (**f**) represent the 95% Confidence Interval (CI). *P < 0.05; **P < 0.01; ***P < 0.001; N.S., not significant. We verified that the Experiment 3 cohort with eye-tracking exhibited the same behavioral signatures of learning as the full cohort. The cohort retained a robust modulation of RTs by syllable position in structured streams (Greenhouse-Geisser corrected *F*(1.76, 67.05) = 86.34, *p* < 0.001, partial η^2^ = 0.69; Supplementary Fig. 3d). Interaction between stream type and position was significant (Greenhouse-Geisser corrected *F*(1.89, 71.92) = 52.12, *p* < 0.001, partial η^2^ = 0.58) and RTs modulation was larger in structure than in random streams(*t*(38) = 9.341, p < 0.001, CI: [0.08, 0.12], Cohen’s *d* = 1.47), further confirming successful statistical learning, with lacking RTs modulation between first and second syllable in random streams (*t*(38) =-1.384, *p* = 0.175, CI: [-13.6, 3.92], Cohen’s *d_z_* = 0.22, *BF*_10_ = 0.42; Supplementary Fig. 3e). Furthermore, the event-driven distortion of subjective time replicated in this group. An LMM analysis confirmed a significant interaction between session (structured vs. random) and condition (within-words vs. between-words; *β* = –0.004, *t*(11190) = –3.580, *p* < 0.001, CI: [-0.007, - 0.002]), driven by only the compression of pauses within words (Estimate =-0.004, *t*(11190) =-4.426, *p* < 0.001, CI: [-0.006,-0.002]) without the expansion of pauses between words (Estimate = 0.0006, *t*(11190) = 0.524, *p* = 0.524, CI: [-0.001, 0.002], *BF*_10_ = 0.18 < 1/3) in the structured session relative to the random session (Supplementary Fig. 3f). Finally, no significant correlation was observed between the RT-based prediction index and the magnitude of the pause-distortion effect (*r* =-0.160, *p* = 0.328, CI: [-0.45, 0.16], *BF*_10_ = 0.20 < 1/3, Supplementary Fig. 3g). These findings confirm that the eye-tracking cohort in Experiment 3 remains behaviorally representative, and again proved the robustness of the shifts for the temporal distortion when adding semantic meaning. In Experiment 3, FOOOF-corrected power spectra revealed clear peaks at the syllable (2.667 Hz) and word (0.889 Hz) frequencies in structured streams. Paired *t*-tests with four surrounding bins confirmed significant entrainment at both syllable (*t*(38) = 5.548, *p* < 0.001, CI: [0.09, 0.19], Cohen’s *d_z_* = 0.55) and word frequencies (*t*(38) = 6.190, *p* < 0.001, CI: [0.04, 0.08], Cohen’s *d_z_* = 0.26). In random streams, a significant peak emerged only at the syllable frequency (*t*(38) = 5.760, *p* < 0.001, CI: [0.08, 0.17], Cohen’s *d_z_* = 0.53), while the word peak was absent (*t*(38) =-1.346, *p* > 0.05, CI: [-0.03, 0.01], Cohen’s *d_z_* = 0.05, *BF*_10_ = 0.40). A 2×2 ANOVA revealed a significant interaction between frequency and stream type (*F*(1, 38) = 7.793, *p* = 0.008, partial η^2^= 0.17), with a higher relative power for structured streams than random streams at the word frequency (*t*(38) = 5.592, *p* <0.001, CI: [0.04, 0.09], Cohen’s *d* = 0.91) and no significant difference at the syllable frequency (*t*(38) = 0.817, *p* = 0.419, CI: [-0.02, 0.04], *BF*_10_ = 0.24 < 1/3), indicating selective tagging of word-level structure in contexts with events (Supplementary Fig. 3h).

## Notes

### Competing Interest Statement

The authors have declared no competing interest.

